# Human cerebral assembloids as a model to study the interaction of glioblastoma with resident glial cells

**DOI:** 10.64898/2026.09.29.755236

**Authors:** Sofia Kalpazidou, Polyxeni Moysidou-Tsiorva, Buse Özbaykent, Ecem Balcioglu, Maria. L. Richter, Elijah R. Shelton, Michael Frischmann, Jovana Spalevic, Patrick N. Harter, Claire Delbridge, Arthur Wagner, Maria Goldberg, Chiara Negwer, Bernhard Meyer, Jürgen Schlegel, Friedhelm Serwane, Julia Schneider, David M. Bühler, Rainer Glass, Milos Ninkovic, Veronica Pravata, Ryszard Wimmer, Ikka Miinalainen, Silvia Cappello, Stefan H. Stricker, Anna Danese, Jovica Ninkovic

**Affiliations:** Chair of Cell Biology, Biomedical Center (BMC), LMU Medizin, LMU Munich, Germany; Institute of Stem Cell Research, Helmholtz Center Munich, German Research Center for Environmental Health, Neuherberg, Germany; Faculty of Physics and Center for NanoScience, Ludwig-Maximilians-Universität in Munich, Germany; Institute of Biophysics, Ulm University, Ulm, Germany; Institute of Neuropathology, LMU Medizin, Ludwig-Maximilians-Universität München, Munich, Germany; German Cancer Consortium (DKTK) partner site Munich, Munich, Germany and German Cancer Research Centre (DKFZ), Heidelberg, Germany; Comprehensive Cancer Center (CCC) Munich, University Hospital, Ludwig-Maximilians University München, Munich, Germany; Bavarian Cancer Research Center (BZKF), Munich, Germany; Department of Pathology, Technical University Munich, Munich, Germany; Deptartment of Neurosurgery, Klinikum Rechts der Isar, Munich, Germany; Department of Neuropathology, Institute of Pathology, School of Medicine and Health, Technical University Munich, 81675 Munich, Germany; Exercise Biology, Department of Sport and Health Sciences, TUM School of Medicine and Health, Technical University Munich, 80809 Munich, Germany; Excellence Cluster for Systems Neurology (SyNergy), Munich, Germany; Department of Neurosurgery, Neurosurgical Research, LMU University Hospital, LMU Munich; Chair of Physiological Genomics, Biomedical Center (BMC), LMU Medizin, LMU Munich, Germany; Max Planck Institute of Psychiatry, Munich, Germany; Biocenter Oulu, University of Oulu, P.O. Box 8000, FI-90014 Oulu, Finland; Institute of Stem Cell Research, Helmholtz Center Munich, Grosshadernerstr.9, Planegg-Martinsried 82152, Germany; IRU Central Nervous System Regeneration, Helmholtz Center Munich, Grosshadernerstr.9, Planegg-Martinsried 82152, Germany

## Abstract

Glioblastoma (GB) is a highly invasive brain cancer characterized by extensive peritumoral glial reactivity. Although GB infiltration and recurrence remain major therapeutic challenges, the contribution of GB-induced gliosis to tumor progression is poorly understood. To investigate glia–GB interactions, we developed human iPSC-derived cerebral forebrain assembloids containing innate microglia and macroglia and transplanted them with GB cells. The model recapitulates key tumor features observed in patients, including distinct tumor regions, reactive gliosis, and altered tissue stiffness. Single-cell RNA sequencing revealed an unexpected enrichment of extracellular vesicles (EVs) in peritumoral reactive glia, which we validated in patient samples. EV inhibition impaired formation of the peritumoral niche, whereas EVs from GB-containing assembloids induced glial reactivity in GB-free assembloids. Furthermore, the modularity of our system enables incorporation of additional niche components, such as monocytes, allowing investigation of the monocyte–glia–tumor axis and providing a powerful platform to dissect mechanisms regulating the GB microenvironment.

## Introduction

Glioblastoma (GB) is the most aggressive, invasive, and lethal brain tumor in adults, classified as a Grade 4 malignancy by the World Health Organization ^1^. Even with extensive treatment, including surgical resection followed by multiple rounds of chemotherapy and radiotherapy, the median survival time from diagnosis is only 14 months ^2^, while median survival time of untreated patients is about 3 months post-diagnosis. Despite significant advances in the field of tumour biology and novel treatment approaches, the overall prognosis remains dismal, with only 5% of patients surviving up to 5 years after diagnosis ^1,2^. Tumor relapse plays a major role in this high mortality rate, as over 90% of patients experience recurrence within a few months after treatment ^3–5^. Notably, recurrent GB foci are primarily located near the resection margins, defined by the peritumoral area (PA) ^3–5^. Therefore, understanding the interactions between tumor cells and brain-resident cells in the PA is critical for identifying patients who might benefit from supramarginal resection and ultimately improving treatment outcomes ^6,7^.

The PA is a complex microenvironment induced by GB, characterized by dynamic interactions among various cell types and the extracellular matrix ^8^. Notably, glial cells in the PA, particularly astrocytes and microglia, exhibit marked differences from their counterparts in the healthy brain ^9^. These differences, which include alterations in cellular morphology, abundance, transcriptomic profiles and secretome, are collectively referred to as reactive gliosis in the PA and can regulate GB development and progression ^10^. Indeed, tumor-associated microglia and macrophages (TAMs) constitute the largest fraction of non-malignant cells in the PA ^11^. Moreover, specific microglial states that suppress adaptive immune responses have been observed in the glioma microenvironment, as in other tumor types ^12^. Increased numbers of TAMs are associated with higher glioma grades and poorer prognoses ^13–15^, while reducing TAMs through blockade of the colony-stimulating factor-1 receptor (CSF-1R) has led to transient regression of GBs in preclinical models ^16^. Similarly, several reports indicate that reactive astrocytes are frequently and closely associated with glioma cells in the central nervous system (CNS) ^17–19^, influencing GB progression, aggressiveness, and patient survival through the release of various cytokines ^20^. Furthermore, astrocyte-mediated alterations in ion concentrations can lead to a loss of epithelial polarization and initiate the epithelial-to-mesenchymal transition, thereby contributing to the invasiveness of GBs ^21^.

Reactive glial cells are commonly found in various pathologies of the CNS, where they form a glia-enriched area that isolates affected tissue from healthy brain parenchyma ^22^. However, activated glial cells can dramatically modify the extracellular environment, sometimes leading to fibrotic scar formation, as observed following spinal cord injury ^22^. Interestingly, recent evidence suggests that a fibrotic area is built in the PA and that inhibition of fibrosis significantly reduces tumor recurrence ^16^. This finding indicates a direct link between alternations in the PA and tumor progression and underscores the need for further investigation into these interactions. Moreover, changes in the stiffness of the extracellular matrix during scar formation can facilitate cell migration and promote tumor invasion, further highlighting the complexity of the PA and revealing diverse mechanisms regulating GB development and progression^23^.

In summary, the PA is shaped by complex interactions among GB cells, neural and immune cells, and various components of the extracellular matrix. This complexity makes it challenging to decipher the specific roles and contributions of each cell type to GB progression and recurrence, as well as to assess their suitability for targeted therapies ^24^. To address this challenge, we developed a human, modular, simplified 3D model to study the emergence and function of the PA in GB progression. This model relies on transplanting either primary, patient isolated GB cells or GB cell lines into human cerebral assembloids (hc-assembloids) derived from induced pluripotent stem cells (iPSCs). In contrast to previous GB organoid models ^25^, the PA in our system develops locally as the tumor progresses, faithfully recapitulating key histological, transcriptomic, and biophysical properties of human GB and its peritumoral microenvironment. Moreover, we demonstrated the modularity of the model by co-culturing tumor-containing human cerebral assembloids with iPSC-derived monocytes, which invaded the PA and GB in a manner consistent with patient observations. This platform provides a unique opportunity to study the interactions between GB and its PA, paving the way for dissecting the roles of different cellular populations and their extracellular matrix in GB progression and recurrence, and enabling high-throughput drug testing ^26^.

## Results

### Unravelling the Cellular Heterogeneity of Human Cerebral Assembloids

To model the interaction between glioblastoma (GB) and surrounding glial cells, we aimed to develop a system incorporating both neurons and glial cells, which are essential components of the peritumoral area (PA). To achieve this, we combined three established protocols ^27–29^ (**Fig.1a**) to generate human cerebral assembloids (hc-assembloids) containing excitatory and inhibitory neurons, as well as glial cells, including microglia (**Fig.1c-j**). The core of our combined protocol involved establishing ventral and dorsal identities using small molecule treatments at the embryoid body (EB) stage (days 5-12, **Fig. 1b**) ^27,29^, along with neuroectoderm induction using a reduced concentration of heparin. This lower concentration enabled neuroectoderm induction, characterized by a distinct translucent, radially organized zone (**Fig. 1b′**), while it allowed some cells to enter the mesodermal lineage and generate microglia ^28^. On day 13, patterned dorsal and ventral EBs, were fused and embedded in Matrigel matrix droplets. From days 13-17, these hc-assembloids were maintained without agitation in Improved Differentiation Medium without vitamin A (IDM-A) to support further neuroepithelial bud expansion. On day 17, hc-assembloids were transferred to IDM+A medium and cultured with agitation (55 rpm orbital shaker), with media changes every three days. By day 30 of culture, expanded neuroepithelial buds were evident (**Fig. 1b′′-b′′′**) and the culture was continued until day 132. This extended culture period aligns with previous reports suggesting that glial cell development requires approximately four months of organoid maturation ^30^.

**Figure 1.**
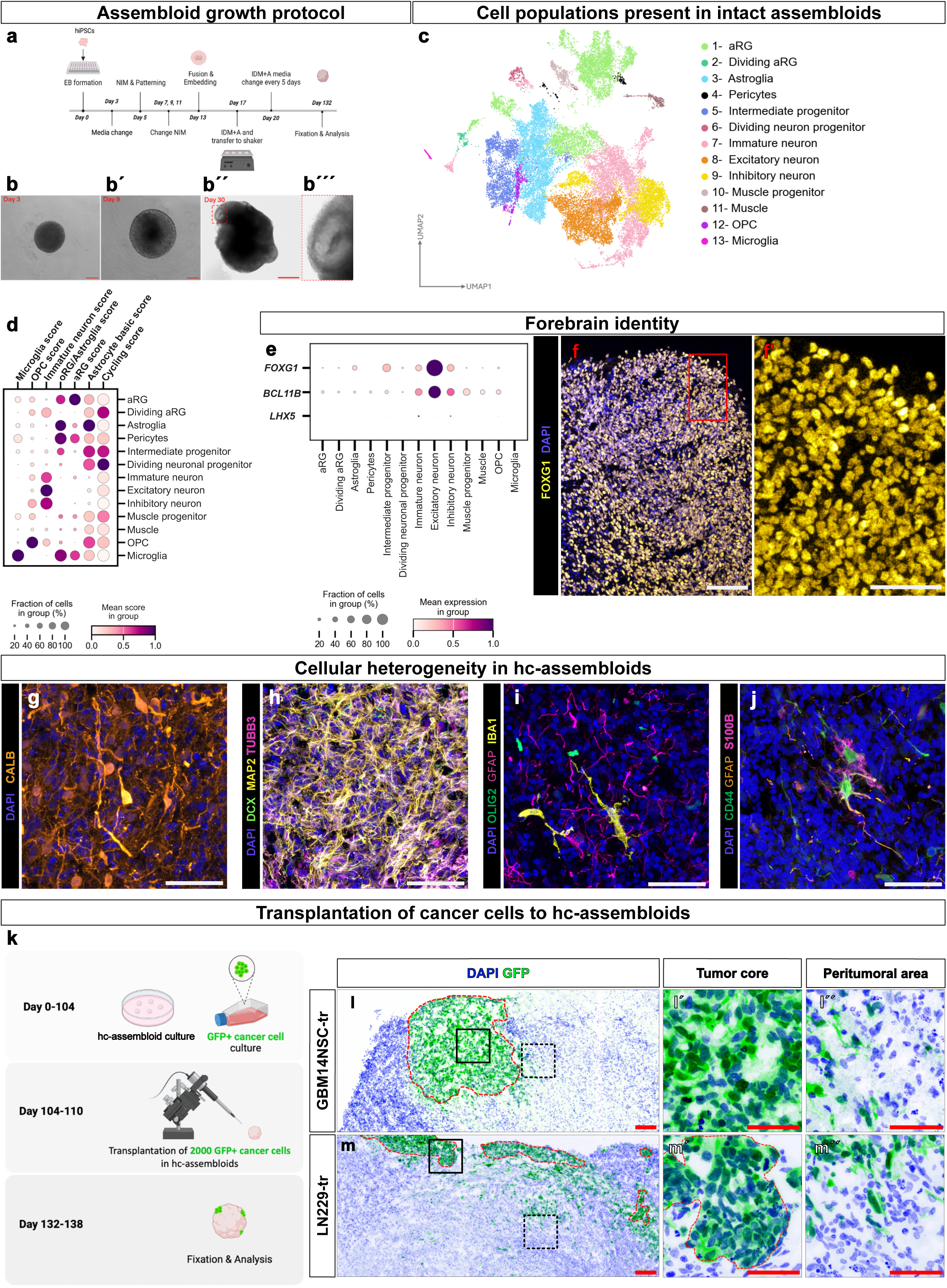
Cellular heterogeneity in hc-assembloids and establishment of the GB-hc-assembloid platform. (a) Schematic representation of the adapted protocol and timeline for generating hc-assembloids from hiPSCs, depicting the formation of a distinct neuroectoderm layer and subsequent expansion of neuroepithelial buds. (b-b′′′) Phase-contrast images illustrating the morphological progression of hc-assembloids at 3 (b), 9 (b′) and 30 (b′′, b′′′) days of culturing as described in Lancaster et al., 2014 ^27^. b′′′ highlights neural rosettes structures at day 30, indicative of the self-organizing 3D structure. (c) UMAP plot showing the cellular clusters present in intact 130 days old hc-assembloids (26039 cells). Clusters are color-coded and annotated according to their transcriptional identities. (d) Dotplot depicting the relative expression of signature scores identifying cell types across the annotated clusters. Dot size represents the fraction of cells within each cluster and color intensity indicates mean score in group. (e) Dotplot depicting the expression of forebrain (FOXG1 and BCL11B), and midbrain/hindbrain (LHX5) identity markers, in 130 days old hc-assembloids. Dot size represents the fraction of cells within each cluster and color intensity indicates mean expression in group. (f) Micrographs depicting expression of FOXG1 in 130 days old hc-assembloids. The red box indicates the magnified area in f′. (g-j) Micrographs depicting the expression of interneurons (g), neuronal cells (h), and glial lineage (I, j) markers. (k) Scheme depicting experimental workflow for the transplantation of GB cells to hc-assembloids. (l, m′′) Micrographs depicting GFP-positive GB cells 3,5 weeks after the transplantation to hc-assembloids for GBM14NSC (l, ĺ, ĺ′) and LN229 (m, m′, m′′) cells. l and m are overview images of tumor areas in hc-assembloids (red dashed lines). ĺ, m′ are magnified images of tumor core areas (black box) and ĺ′, m′′ are magnified images of peritumoral areas (black dashed box) with GFP-positive GB cells and GFP-negative host cells. Scale bars: 100 μm in f, l, m and 50 μm in f’, g, h, I, j, ĺ, ĺ′, m′, m′′. Abbreviations: EB-Embryoid Bodies, NIM-Neuronal Induction Medium, IDM+A-improved differentiation media with vitamin A, Astrocyte, OPC-Oligodendrocyte Progenitor Cells, GFP, hc-human cerebral, tr-transplanted

To assess the cellular heterogeneity of hc-assembloids at day 132, we performed single-cell RNA sequencing (scRNA-seq) using cells isolated from 9 intact hc-assembloids (derived from 3 independent batches). To ensure an unbiased evaluation, we employed a recently developed protocol that does not selectively enrich specific cellular populations ^31,32^. Cellular heterogeneity was visualized using Uniform Manifold Approximation and Projection (UMAP) (**Fig. 1c, Fig. S1a**), revealing 36 cellular clusters (**Fig. S1a**). Using previously published human organoid datasets as reference ^33^, we identified multiple cell types (**Fig. S1a**) including apical radial glia (aRG), homeostatic astrocytes, oligodendrocyte progenitor cells (OPCs), interneuron progenitors, and neurons/neuronal progenitors along with mesenchymal lineage-(microglia and muscle lineage) and neural crest-derived (pericytes) cells (**Fig. 1c, d**). Neuronal, astrocytic, and neural progenitor clusters exhibited expression of *FOXG1* and *BCL11B* while showing absence or low levels of *LHX5* (**Fig. 1e, f**), consistent with their telencephalic identity ^34^. Notably, mesenchymal clusters lacked telencephalic *FOXG1* expression (**Fig. 1e**). To assess dorsal and ventral telencephalic identities, we generated a gene expression identity scores (**Table S1**) based on previously reported gene expression data ^29,35^. This analysis confirmed the presence of both dorsal and ventral identity cells in our model (**Fig. S1b,c**). Consistent with the presence of ventral identity, we observed GB-relevant interneuron populations, such as calbindin-positive interneurons (**Fig. 1g**), alongside general neurons and neural progenitors (**Fig. 1h**).

Since our goal was to study glial cell interactions with GBs, we next examined the presence of, oligodendroglia, astroglia and microglia in hc-assembloids at day 132 (**Fig. 1i, j** and **Fig. S1d-n**). To analyze oligodendrocyte progenitor cell (OPC) emergence, we generated an OPC expression score (**Table S2**) based on 19 human OPC-specific markers ^36–38^. The OPC expression score revealed the presence of an OPC cluster, namely cluster 35 (**Fig. 1d**), including strong expression of the oligodendroglial lineage marker *OLIG2.* OLIG2-positive cells were also detected in organoids using Immunofluorescence (IF) analysis (**Fig. 1i**).

We identified several cellular clusters expressing common astrocyte markers such as *GFAP*, *S100B*, *ALDH1L1*, *SOX9*, *AQP4*, *GJA1*, and *CD44*. Some of these exhibited gene expression patterns indicative of apical radial glia (aRG) ^30,33^ (**Table S3**, **Suppl. Fig. S1a**), while several clusters showed expression patterns typical for more mature astrocytic states (**Fig. S1d**). Amongst those clusters we identified the gliogenic progenitors, mature homeostatic astrocytes and ependymal cells (**Fig. S1 e-h**). Correspondingly, IF analysis detected non-radial GFAP-, CD44-, and S100B-positive parenchymal cells (**Fig. 1j**), suggesting the presence of post-mitotic astrocytes in our organoid system, further validated by the lack of proliferation in these clusters (**Fig. S2n, o**).

We generated a microglia expression score using 26 signature genes previously identified as microglia-specific markers (**Table S4**) ^28,39^. This score was exclusively detected in cluster 36 (**Fig. S1a** and **Fig. 1d**). Furthermore, *PTPRC* expression, indicative of mononuclear lineage cells, was specifically observed in microglial cluster (**Fig. S1i**). Flow cytometry (FACS) analysis confirmed CD45 antigen expression on 0.25% of hc-assembloid cells (**Fig. S1j, k**), consistent with scRNA-seq analysis showing that microglial cluster comprised ∼0.3% of all identified cells. Importantly, CD45 levels in hc-assembloid-derived microglia was lower than in peripheral blood monocytes (PBMCs) (**Fig. S1l, m**), aligning with their microglial identity. Similarly, the microglial marker *AIF1* was expressed in 90% of cluster 35 cells (**Fig. S1i**). IF analysis against IBA1 (encoded by *AIF1*) revealed microglial cells with a ramified morphology in 132-day-old hc-assembloids (**Fig. 1i**), indicative of a homeostatic state ^40^. To further assess microglial maturation, we examined “sensome” gene expression, which has been reported to increase only 24 weeks post-transplantation of organoids into rat brains ^39^. Indeed, in our system, microglial cells showed strong enrichment of multiple “sensome” genes (**Fig. S1n**) without any embedding into an animal model system.

In conclusion, our single-cell transcriptomic and IHC data indicate that our model contains all essential glial populations required to study human glial interactions with GB.

### Glioblastoma Cells Form Tumor Foci Following Transplantation into hc-Assembloids

To investigate the interaction of GB cells with the glial microenvironment, we used two GB cell lines, **GBM14NSC** ^41^ and **LN229** ^42^. Prior to transplantation, we performed single-cell RNA sequencing to characterize the transcriptional profiles of the parental cell lines (**Fig. S2a, b**). Consistent with their high proliferative capacity, cells from both lines exhibited elevated cell-cycle scores (**Fig. S2c**). Cell-cycle scores were calculated using the KEGG Cell Cycle gene set obtained through the Harmonizome platform ^43,44^. Despite their similar proliferative states, GBM14NSC and LN229 cells (**Fig. S2a, c**) clustered separately (**Fig. S2a**), with only a small number of clusters containing cells from both lines (**Fig. S2b-d**).

To define the molecular subtype of each cell line, we scored cells using established GB state-specific gene signatures ^45^. LN229 cells predominantly exhibited a MES1 mesenchymal-like state (**Fig. S2e**), whereas GBM14NSC cells primarily displayed astrocyte-like (AC-like) and oligodendrocyte progenitor-like (OPC-like) states (**Fig. S2e**). In addition, one GBM14NSC cluster exhibited a neural progenitor cell-like (NPC-like) state (**Fig. S2e**). Interestingly, the mixed clusters containing both LN229 and GBM14NSC cells predominantly displayed either AC-like or OPC-like transcriptional programs (**Fig. S2e)**. Together, these findings demonstrate that the two GB models represent multiple, distinct transcriptional states, enabling the investigation of both shared and subtype-specific interactions with the surrounding brain tissue. To assess these interactions in a human brain-like environment, approximately 2,000 GFP-labeled GB cells were transplanted into 104-110-day-old hc-assembloids and analyzed during 3.5 weeks after transplantation (**Fig. 1k**). As early as 5 days post-transplantation (dpt), scattered GFP-positive cells were detected within the hc-assembloid tissue for both GB lines (**Fig. S2f, i**). Over time, these cells progressively invaded the surrounding tissue and formed well-defined tumor foci (**Fig. S2g-k**). Histological analysis of transplanted hc-assembloids (n = 15 hc-assembloids from 10 independent differentiations) identified three distinct anatomical regions **(Fig. 1l-m′′)**: (i) the **tumor core (TC)**, composed exclusively of GFP-positive GB cells **(Fig. 1l,m,l′,m′)**; (ii) the **peritumoral area (PA)**, containing both GFP-positive GB cells and GFP-negative resident hc-assembloid cells (**Fig. 1l,m,ĺ′,m′′′**); and (iii) the **tumor-free region (TF)**, consisting exclusively of resident hc-assembloid cells (**Fig. 1l,m**). Notably, the PA exhibited poorly defined borders, with numerous GB cells infiltrating and intermingling with the surrounding healthy tissue (**Fig. 1l, m**), closely resembling one of the defining histopathological features of human glioblastoma and a major obstacle to successful surgical resection and therapy ^5,46^. Live imaging of 300 μm-hc-assembloid-slices containing GB foci further revealed highly dynamic interactions between invading GB cells and surrounding non-neoplastic host cells (**Video S1, S2 and S3**).

Moreover, long-term imaging revealed migration of GB cells through the PA, consistent with their invasive capacity (**Video S3**). Importantly, the architecture of the PA differed markedly between the two tumor models. LN229 tumors developed a substantially broader PA containing numerous individually invading GFP-positive GB cells, whereas GBM14NSC tumors displayed a more compact invasive front (**Fig. 1l, m**). This phenotype is consistent with the predominantly MES1 transcriptional state of LN229 cells, which has been associated with enhanced migratory and invasive behavior including PA infiltration ^45,47^

Together, these findings demonstrate that our hc-assembloid model faithfully recapitulates key structural features of the glioblastoma peritumoral brain zone, including the coexistence of infiltrating tumor cells with resident glial and neuronal populations, closely resembling the pathological organization observed in patients ^48^.

#### The hc-Assembloid Model Recapitulates GB Subtype-Specific Features

To determine whether and how the hc-assembloid-specific environment influences GB cells, we compared gene expression in two GB cell lines, before transplantation, cultured as GB cell cultures (GB-C), and 3.5 weeks after transplantation into hc-assembloids (GB-A), (**Fig. S2l**). Following data integration and batch correction, LN229 cells within the hc-assembloids were identified post hoc based on GFP expression, whereas GBM14NSC cells were identified based on high expression of OS9, a commonly used GB marker ^49^ (**Fig. S2m**). Non-neoplastic cells did not express either of these markers, and their identities were determined using human organoid datasets as a reference ^33^ (**Fig. S2n**).

We first extracted the GB cells from the full dataset (**Fig. S2m**) and further subclustered them (**Fig. 2a**). This analysis revealed that the hc-assembloid environment induced transcriptional changes in GB cells, resulting in the emergence of clusters that were detected only after transplantation (**Fig. 2b, Fig. S3a-c**). Importantly, however, transplantation did not alter GB subtype identity. LN229 cells retained a predominantly MES1 phenotype, whereas GBM14NSC cells expressed markers of AC-like and OPC-like states in both GB-C and GB-A clusters (**Fig. 2c**). Moreover, mixed clusters containing both GBM14NSC and LN229 cells displayed an OPC-like phenotype, regardless of whether they contained GB-C or GB-A cells. The hc-assembloid environment appeared to exert a stronger influence on LN229 cells, as we observed almost no mixing of the LN229-C and LN229-A conditions within the same clusters. This was evident both in MES1 clusters containing LN229-A cells, including clusters 20 and 8, and in clusters containing LN229-C cells, including clusters 7, 13, and 19 (**Fig. 2b**).

**Figure 2.**
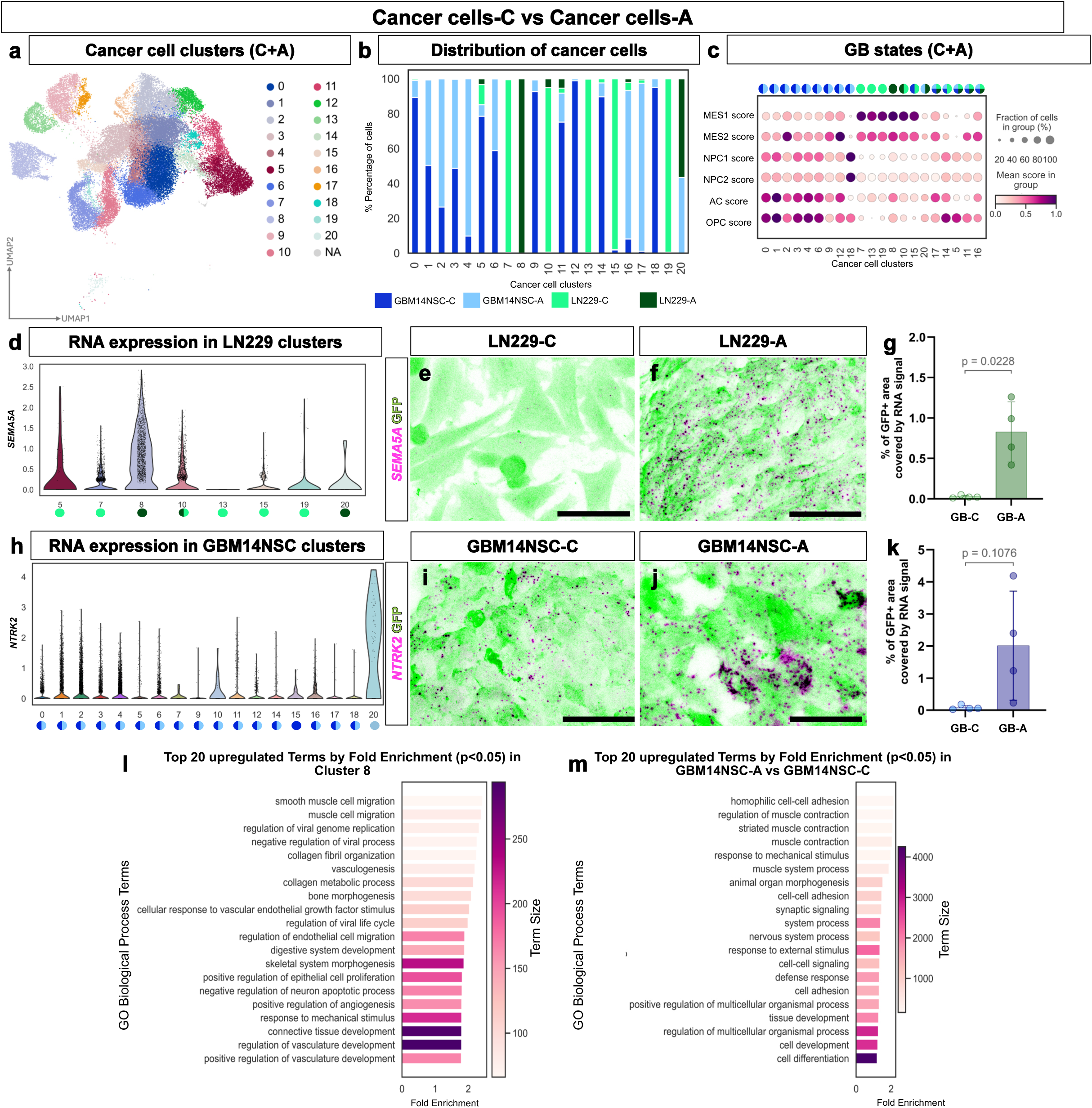
Characterization of cancer cells before and after transplantation in hc-assembloids. (a) UMAP plot depicting cancer cell clusters of GBM14NSC and LN229 cells before (GB-C) and after (GB-A) transplantation. (b) Bar plot displaying the fraction of cells from each experimental condition within identified GB clusters. (c) Dot plot demonstrating GB cellular state signatures across cancer cell clusters. MES1, MES2, NPC1, NPC2, AC, and OPC scores were calculated for each cluster as described in Neftel et al.^45^. Contribution of experimental conditions to the identified clusters is color-coded above the dotplot. Dot size represents the fraction of cells within each group and color intensity indicates mean score withing the group. (d) Violin plot showing expression levels of *SEMA5A* across LN229 clusters. (e, f) Micrographs depicting RNA scope signal for *SEMA5A* RNA in LN229-C (e) and LN229-A (f) cancer cells. (g) Dot plot showing the percentage of the GFP-positive area covered by *SEMA5A* RNA signal in GB-C and GB-A condition. Data are shown as mean ± standard deviation (SD). Each dot represents biologically independent samples (passages for GB-C and transplanted hc-assembloids from two different batches for GB-A). Normality was assessed using the Shapiro-Wilk test, followed by an unpaired two-tailed t-test, p < 0.05. (h) Violin plot showing expression levels of *NTRK2* across GBM14NSC clusters. (I, j) Micrographs depicting RNA scope signal for *NTRK2* RNA in GBM14NSC-C (i) and GBM14NSC-A (j) cancer cells. (k) Dot plot showing the percentage of the GFP-positive area covered by *NTRK2* RNA signal in GB-C and GB-A condition. Data are shown as mean ± SD. Each dot represents biologically independent samples (passages for GB-C and transplanted hc-assembloids from two different batches for GB-A). Normality was assessed using the Shapiro-Wilk test, followed by an unpaired two-tailed t-test, statistical significance was defined as p<0.05. (I, m). Bar plots showing the top 20 significantly enriched Gene Ontology (GO) Biological Process terms (p < 0.05) in the genes upregulated in cluster 8 compared to all other LN229 clusters (l) and in GB-A conditions compared to GB-C conditions for GBM14NSC (m), ranked by fold enrichment. Color intensity indicates the number of genes associated with each GO term (term size). Scale bars: 50 μm in e, f, i, j. Abbreviations: -C-culture, -A-assembloid, MES-Mesenchymal, NPC-Neural Progenitor Cells, AC-Astrocyte, OPC-Oligodendrocyte Progenitor Cells, GFP-Green Fluorescent Protein, logFC-log fold change, GO-Gene Ontology

Because GB cells were transplanted in hc-assembloid medium, we next examined whether the observed transcriptional changes could have been induced by the culture medium independently of transplantation. To address this possibility, we included an additional condition in the third biological replicate in which GBM14NSC cells were cultured in organoid medium (GBM14NSC-C + IDM+A medium, **Fig. S3l-n**; see Materials and Methods). Following data integration, we compared the three datasets using cosine similarity, Pearson correlation coefficients, and nearest-neighbour (NN) mixing analysis. The NN mixing analysis showed that GBM14NSC cells cultured in organoid medium more closely resembled GBM14NSC cells maintained under standard culture conditions than transplanted GBM14NSC cells across all replicates (**Fig. S3n**; **Suppl. Information Table 5**). These findings exclude the possibility that the transplantation-induced transcriptional changes were attributable mainly to differences in culture conditions.

To further investigate the basis of this transcriptional separation, we performed differential gene expression analysis between cluster 8, which contained only LN229-A cells, and all other LN229 clusters. This analysis identified 3119 significantly regulated genes (q value < 0.05; log_2_FC > 0.6 for upregulated genes and log_2_FC < −0.6 for downregulated genes; mean expression > 0.2). To validate these findings, we selected two upregulated genes enriched in either cluster 8, which displayed a MES1 phenotype, or cluster 20, which displayed an OPC progenitor-like phenotype (**Fig. 2d, Fig. S3d**). The selected genes, *SEMA5A* and *NR4A1*, are relevant to glioblastoma biology, contributing to tumor progession, invasiveness and survival ^50,51^. We analyzed their expression in the GB-A and GB-C conditions using the orthogonal RNAscope method. This analysis confirmed the significant upregulation of the selected genes post-transplantation (**Fig. 2e-g, Fig. S3e-g**). Gene Ontology (GO) analysis of genes upregulated in cluster 8, which contained transplanted LN229 cells, revealed enrichment of biological processes associated with “vasculogenesis,” “positive regulation of angiogenesis,”, “regulation of vasculature development” and “smooth muscle cell migration” (**Fig. 2l**). Because our hc-assembloid model lacks intrinsic vasculature, these findings likely reflect an inherent capacity of LN229 cells to modify their microenvironment and promote vascular remodelling, a well-documented feature of GB cells in vivo ^52^. We also observed enrichment of processes related to collagen fibril organization and the response to mechanical stimuli (**Fig. 2l**), suggesting a specific interaction between GB cells and their environment following transplantation.

Unlike LN229 cells, which were predominantly segregated into two transplantation-specific clusters, GBM14NSC-A cells were distributed across clusters containing both GB-A and GB-C cells (**Fig. 2c**). We therefore compared the transcriptomic profiles of GB-A and GB-C cells within these mixed clusters to determine how the hc-assembloid environment influences the GBM14NSC state. Differential gene expression analysis identified 718 significantly regulated genes (q value < 0.05; log_2_FC > 0.6 for upregulated genes and log_2_FC < −0.6 for downregulated genes; mean expression > 0.1), including the cancer-relevant genes *NTRK2* and *GRID2* ^53,54^ (**Fig. 2h, Fig. S3h**). Their differential expression was validated using RNAscope (**Fig. 2i-k, Fig. S3i-k**). GO analysis of genes upregulated in the GB-A condition revealed enrichment of biological processes related to synaptic signalling and nervous system function (**Fig. 2m**). These findings are consistent with previously reported glioblastoma– neuron networks ^55^ and may indicate a stronger commitment of GBM14NSC cells to a proneural-like state following transplantation. Moreover, enrichment of GO terms related to cell-cell adhesion, cell-cell signalling, responses to mechanical stimuli, and responses to external stimuli (**Fig. 2m**) indicates extensive interactions between GB-A cells and the surrounding environment.

Transcriptomic analyses of both LN229 and GBM14NSC tumors revealed upregulation of pathways associated with responses to mechanical stimuli, suggesting that biomechanical cues may contribute to tumor adaptation within the PA. The viscoelastic properties of the PA, particularly extracellular matrix (ECM) stiffness, have been implicated in tumorigenesis, invasion, metabolism, and immune evasion ^56–58^. To determine whether our model recapitulates these biomechanical features, we quantified tissue stiffness in the tumor core (TC) and GB cells free (TF) area using the ferrofluidic droplet method ^59,60^. Briefly, ferrofluidic droplets were microinjected into either the TC or TF regions of hc-assembloids (**Fig. S3o**). All cells were labelled with CellBrite Steady membrane stain to facilitate visualization of the tissue architecture (**Fig. S3p, p′**). A controlled magnetic field was then applied to induce local tissue deformation (**Video S4**), and tissue strain was quantified by monitoring the temporal evolution of the droplet–tissue interface. The effective Young’s modulus was calculated by fitting a linear viscoelastic model to the resulting creep curves. These measurements demonstrated that the TC generated by LN229 tumors was significantly stiffer than the corresponding TF region (**Fig. S3q**). Notably, the Young’s modulus of the LN229 TC was approximately 1.6 kPa, closely matching the stiffness of 1.32 kPa measured in human glioblastomas by magnetic resonance elastography ^61^. In contrast, hc-assembloids transplanted with the OPC progenitor-like GBM14NSC line exhibited substantially lower tissue stiffness than both the LN229 TC and the corresponding TF region (**Fig. S3q**). These results indicate that our model captures biomechanical differences between glioblastoma subtypes. They are also consistent with clinical observations associating reduced tumor stiffness with less aggressive disease and with the OPC progenitor-like GB subtype. Whether brain tumor cells preferentially migrate towards stiffer or softer regions is an active field of research ^62^. Our platform will allow the quantification of brain cancer biomechanics in a physiological relevant human system.

In summary, our findings suggest that transplantation into hc-assembloids does not alter GB subtype identity but instead reinforces the original subtype-specific characteristics and promotes interactions with the surrounding non-neoplastic hc-assembloid tissue. Moreover, the system reproduces clinically measured in vivo tumor stiffness, further supporting its suitability as a model for investigating interactions between glial cells and GB cells.

#### GBs Induce Reactive Gliosis in hc-Assembloids

Our data suggest that GBs transplanted into hc-assembloids recapitulate many features of their in vivo counterparts. We investigated whether they also induce glial reactivity resembling that observed in GB patients ^63^. To this end, we transplanted 2,000 GB cells into 104-to 110-day-old hc-assembloids and analyzed their effects on the two major glial populations present in the model: hc-assembloid-derived astrocytes and microglia.

We first assessed the spatial distribution of reactive astrocytes by immunohistochemistry (**Fig. 3**). In intact hc-assembloids, CD44-positive astrocytes were evenly distributed and exhibited only minimal GFAP expression, consistent with a homeostatic state (**Fig. 3a, a’, d**). In contrast, transplantation of either GBM14NSC or LN229 cells induced a marked upregulation of GFAP in astrocytes located adjacent to the tumor bulk (**Fig. 3b, b’, c, c’**). The induced GFAP signal co-localized with CD44, confirming the astrocytic identity of these reactive cells (**Fig. 3e, f**). Notably, individual GFP-positive tumor cells were frequently observed within this reactive astrocytic compartment (red arrowheads in **Fig. 3b’, c’, e-i**), closely resembling the diffuse infiltrative border between tumor and healthy tissue observed in GB patients ^63^. Consistent with patient tissue, GFAP-positive astrocytes within the PA expressed elevated levels of vimentin, a well-established marker of reactive astrogliosis associated with glioblastoma ^64,65^, compared with intact, non-transplanted hc-assembloids (**Fig. 3g–i**). Because ECM remodelling is a characteristic feature of the GB PA ^58^, we next examined LGALS3, a molecule strongly upregulated in proliferative reactive astrocytes ^66^. Immunohistochemistry revealed robust LGALS3 expression in both GBM14NSC- and LN229-transplanted hc-assembloids, predominantly localized to the PA and partially co-localizing with GFAP (**Fig. 3b-c’**). In contrast, intact hc-assembloids exhibited little to no LGALS3 expression, and the limited signal did not colocalize with GFAP, indicating the absence of LGALS3-positive reactive astroglia in intact hc-assembloids (**Fig. 3a, a’**). Importantly, all markers of reactive astrogliosis remained spatially restricted to the PA surrounding both LN229 and GBM14NSC tumors and gradually declined with increasing distance from the tumor bulk (**Fig. 3j–s**).

**Figure 3.**
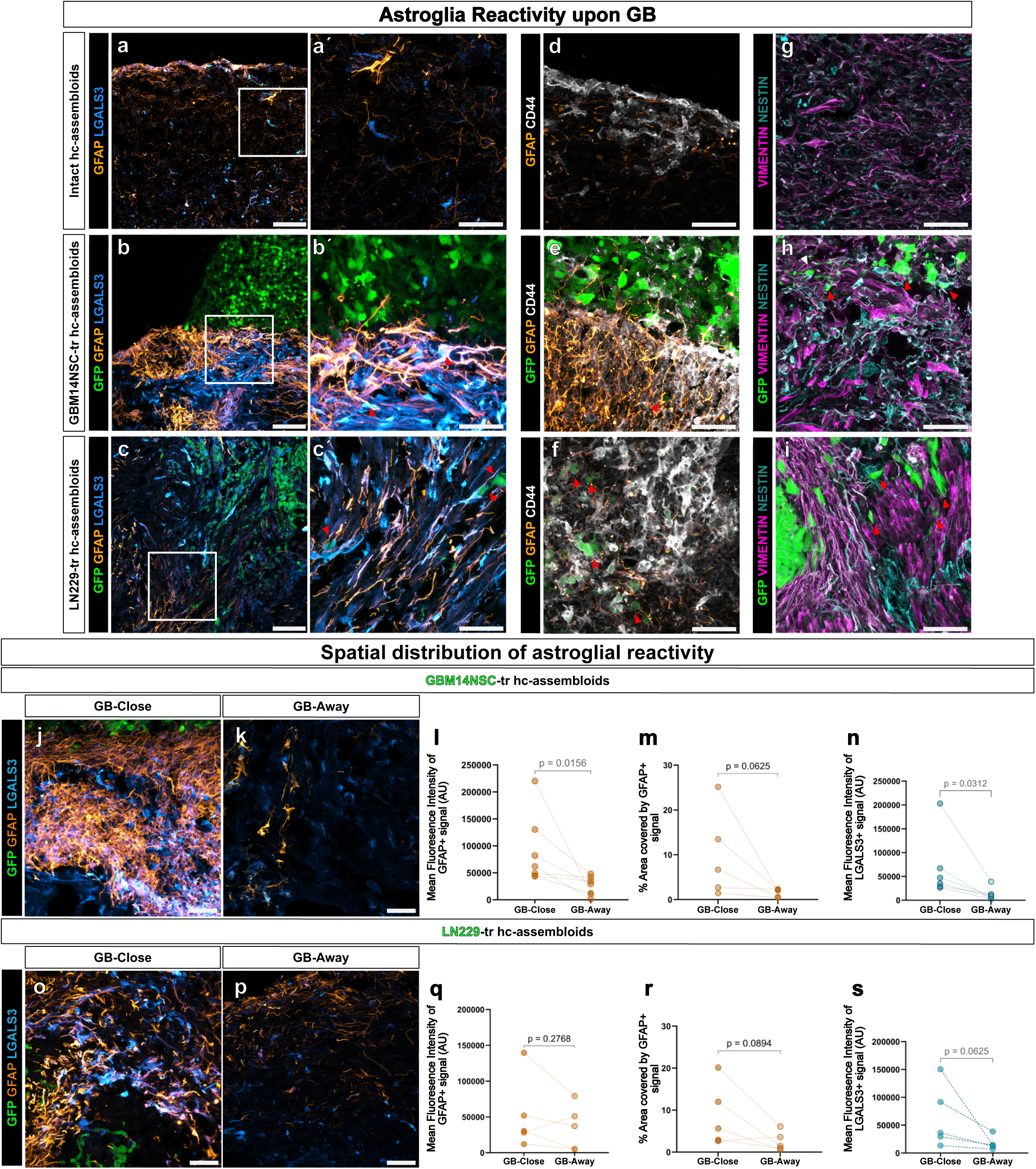
Astrocytic reactivity in GB-containing hc-assembloids. (a-i) Micrographs depicting expression of astrocyte reactivity markers LGALS3 (a, á, b, b′, c, c′), GFAP (a-f), CD44 (d, e, f), Vimentin (g, h, i) and Nestin (g, h, i) in intact (a, á, d, g), GBM14NSC-( b, b′, e, h) and LN229-(c, c′, f, i) transplanted hc-assembloids. (a, b, c) Micrographs are overview images and (á, b′, c′) represent magnified areas (white boxes). (j, k, o, p) Micrographs showing GFAP and LGALS3 immunoreactivity in areas proximal (j, l) and distal (j′, ĺ) to GB niches revealing the spatial pattern of astrocytic reactivity in GBM14NSC-(j, k) and LN229-(o, p) transplanted hc-assembloids. (l, n, q, s) Dot plots depicting GFAP (l, q) and LGALS3 (n, s) mean fluorescence signal intensity in GBM14NSC-(l, n) and LN229-(q, s) transplanted hc-assembloids. (m, r) Dot plots presenting the proportion of area covered by GFAP-signal in GBM14NSC-(m) and LN229-(r) transplanted hc-assembloids. Paired measurements are indicated by dashed lines. Single dots represent individual transplanted hc-assembloids obtained from ≥5 independent batches. Data normality was assessed in all groups with Shapiro-Wilk normality test. Comparisons made using two-tailed paired t-test for (q, r) and two-tailed Wilcoxon matched-pairs signed-rank test for (l, m, n, s). Scale bars: 100 μm in a, b, c and 50 μm in á, b′, c′, d-k, o and p. Abbreviations: GFP-Green Fluorescent Protein, tr-transplanted, hc-human cerebral, GB-Glioblastoma

To determine whether this reactive astrocytic response was specific to GB cells rather than a consequence of transplantation of any proliferative cells, we transplanted iPSC-derived neuronal progenitors (NPCs) generated using previously established differentiation protocols ^67,68^ (**Fig. S4a-c’**). These RFP-expressing NPCs displayed the expected morphology (**Fig. S4b**) and formed a homogeneous PAX6- and TUJ1-positive population (**Fig. S4c, c’**). Because of their substantially lower proliferative capacity compared to GB cells, we transplanted 20,000 progenitor cells per hc-assembloid (ten-fold more than GBM14NSC cells) (**Fig. S4d**). After 3.5 weeks, RFP-positive NPCs had populated the hc-assembloids (**Fig. S4e**) and consisted of both differentiated MAP2-positive neurons and proliferating KI67-positive cells (**Fig. S4f–f′′′**). In contrast to GB transplantation, NPC transplantation induced only minimal GFAP expression and failed to induce LGALS3 expression (**Fig. S4g, g’**). Likewise, neither reactive GFAP/LGALS3-positive astrocytes nor IBA1-positive microglia accumulated around the transplanted NPCs (**Fig. S4i, o**).

We next investigated microglial responses within the PA by immunohistochemistry using IBA1, a well-established marker of microglia/macrophages (**Fig. S5a-e’**). In intact, media-transplanted, and neuronal progenitor-transplanted hc-assembloids, ramified microglia were evenly distributed throughout the tissue (**Fig. S5a-c’**). In contrast, transplantation of either GBM14NSC or LN229 cells resulted in pronounced microglial accumulation within the PA, while microglia remained largely excluded from the tumor bulk (**Fig. S5d-e’**). Peritumoral microglia displayed an amoeboid morphology characteristic of an activated state (**Fig. S5d’, e’**), whereas microglia located further from the tumor retained a more ramified morphology, indicative of reduced activation. Similarly, microglial density was highest in the PA and progressively decreased with increasing distance from the tumor bulk in both GB models (**Fig. S5f-k)**, closely mirroring the spatial distribution observed in GB patient tissue.

We next asked whether the innately developed microglia are required for formation of the PA. To address this question, we generated cerebral cortex organoids ^69^, that contain GFAP-positive astrocytes but lack microglia at 130 days of differentiation (**Fig. S5l**). These organoids were transplanted with 2,000 GBM14NSC cells at day 105 and analyzed 3.5 weeks later. As expected, no IBA1-positive microglia were detected adjacent to the tumor bulk (**Fig. S5m**). Nevertheless, GFAP expression was still elevated in the PA compared with tumor-distant areas (**Fig. S5n-p**), although the response was markedly weaker than that observed in hc-assembloids containing endogenous microglia (**Fig. 3b,j,o**). Strikingly, LGALS3 accumulation was completely absent in the GFAP-positive PA of microglia-deficient organoids and no GFAP- and LGALS3-positive cells were observed (**Fig. S5n,o,q**), indicating a failure to establish the ECM remodelling characteristic of the GB PA ^58^. Because these differences could theoretically arise not only from the absence of microglia but also from differences in regional identity, as cerebral cortex organoids consist exclusively of dorsally patterned cells, we compared tumor-host interactions between dorsal and ventral hc-assembloid populations using the single cell dataset (**Fig. S5r-t**). GB cells and hc-assembloid cells were classified according to dorsal or ventral identity based on gene expression signatures (**Table S1**; **Fig. S5r, t**). Cell-cell interaction strength was calculated as the average expression of ligand-receptor pairs defined with CellPhoneDB ^70^. This analysis revealed no significant differences in the interaction strength between GB cells and dorsal versus ventral host populations (**Fig. S5t**), suggesting that the impaired development of the PA primarily results from the absence of microglia rather than regional identity.

Together, these findings demonstrate that hc-assembloids faithfully recapitulate the formation of a patient-like PA characterized by reactive astrocytes, activated microglia, and extracellular matrix remodelling surrounding both LN229 and GBM14NSC tumors.

#### The hc-Assembloid Model Recapitulates GBM Subtype-Specific Reactive Gliosis

Histological analysis revealed that both GBM14NSC and LN229 tumors induced reactive gliosis in the PA. However, the morphology and characteristics of this response differed markedly between the two GB subtypes. Both GBM14NSC- and LN229-transplanted hc-assembloids displayed a significantly larger GFAP-positive area in the PA compared with regions of the hc-assembloid that were not in contact with GB cells (**Fig. 3m,r**). While GBM14NSC transplantation resulted in strong GFAP expression in reactive astrocytes, LN229 transplantation induced a weaker GFAP signal intensity (**Fig. 3l,q**). The GFAP-positive area surrounding LN229 tumors was more diffuse, with a broader spatial distribution extending into the adjacent tissue, resulting in a less pronounced difference between the PA and the unaffected hc-assembloid tissue (**Fig. 3o,p**). These observations suggested that distinct GB subtypes induce qualitatively different patterns of reactive gliosis.

To characterize these differences in an unbiased manner, we analyzed the non-neoplastic hc-assembloid cells by single-cell RNA sequencing. We extracted all hc-assembloid-derived cells from the integrated dataset (**Fig. S2n**), comprising intact, media-transplanted, GBM14NSC-transplanted, and LN229-transplanted hc-assembloids (**Fig. S6a**). Following data integration and batch correction, we identified 35 cellular clusters using published human organoid datasets as a reference ^33^. To distinguish GB-induced responses from cellular changes caused by the transplantation procedure itself, we first compared intact, media-transplanted, GBM14NSC-transplanted, and LN229-transplanted hc-assembloids. Three complementary metrics were used to assess batch effects and cellular similarity (see Materials and Methods). Although high cosine similarity and Pearson correlation coefficients indicated only minor baseline differences between the conditions, nearest-neighbor (NN) mixing analysis proved considerably more sensitive and detected distinct transcriptional changes in all transplanted conditions relative to intact hc-assembloids. Importantly, the effects induced by GB transplantation were substantially greater than those observed after media transplantation alone (**Fig. S6c**; **Table S6**). Consistent with this analysis, microglia from intact and media-transplanted hc-assembloids exhibited higher expression of the homeostatic markers *TREM2*, *TYROBP*, *P2RY13*, and *SPI1* ^71,72^, than microglia from GB-transplanted hc-assembloids (**Fig. S6f**). Conversely, microglia from tumor-bearing hc-assembloids showed increased expression of activation-associated genes, including *CD68*, *TLR4*, *APOE*, *IL1B*, *CSF1R*, and *MMP9* ^73–75^ (**Fig. S6g**), indicating a transition from a homeostatic to an activated microglial state. Therefore, all subsequent analyses focused on comparisons between GB-transplanted hc-assembloids and media-transplanted controls.

To investigate reactive astrogliosis, we extracted all astroglial cells from the three transplantation conditions (**Fig. 4a**) and reclustered them, identifying 24 astroglial clusters (**Fig. 4b**). While several clusters were composed predominantly of astrocytes from media-transplanted hc-assembloids, others contained cells from all three transplantation conditions. Differential abundance analysis identified five clusters that were significantly enriched following GB transplantation (**Fig. 4d**), which we interpreted as GB-induced astroglial states. Conversely, eleven clusters were significantly depleted in GB-transplanted hc-assembloids (**Fig. 4d**). Among the GB-enriched clusters, cluster 5 contained comparable proportions of GBM14NSC- and LN229-induced astrocytes, whereas two clusters were predominantly enriched following GBM14NSC transplantation and two others following LN229 transplantation (**Fig. 4c**).

**Figure 4.**
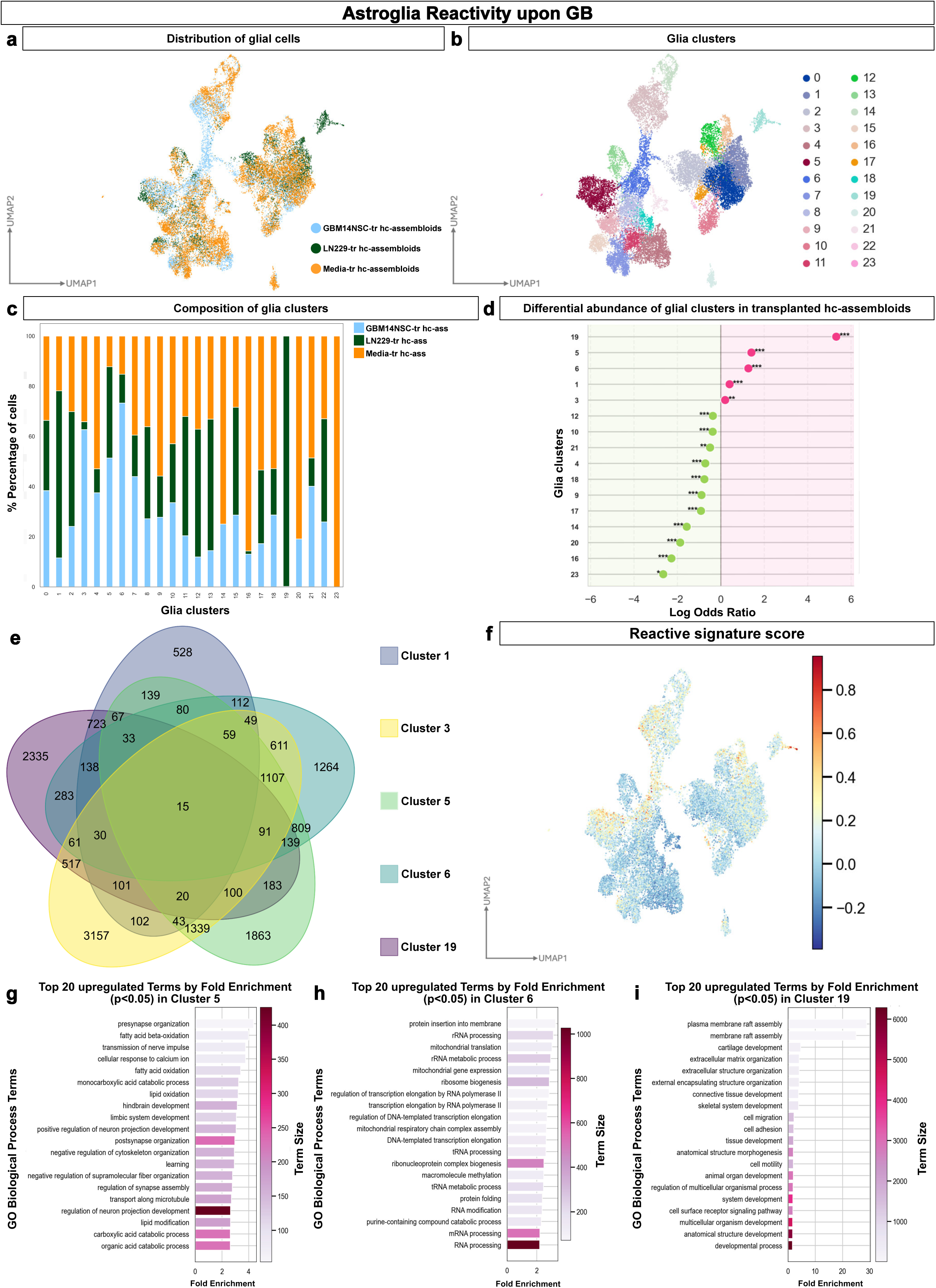
Molecular signatures of reactive astrocytes in GBM14NSC- and LN229-transplanted hc-assembloids. (a) UMAP plot with color coded separation of astroglial cells by experimental condition (21756 cells). (b) UMAP plot showing subclustering of astroglial cells. (c) Bar plot displaying proportion of cells from each experimental condition within identified clusters. (d) Dot plot presenting enrichment of glial clusters measured by log odds ratio in transplanted hc-assembloids compared to control media transplanted hc-assembloids. Positive values denote clusters significantly enriched following transplantation, whereas negative values clusters significantly depleted upon transplantation. Statistical significance is calculated with Wald z-test with Benjamin-Hochberg correction; the significance is indicated by asterisks (*p < 0.05, **p < 0.01, ***p < 0.001) (e) Venn diagram illustrating the intersection of differentially expressed genes between glial cell clusters enriched after GB cell transplantation and all other glial clusters. (f) UMAP depicting the expression score for the reactive signature derived from the 15 genes shared among the GB-enriched glial clusters. (g, h, i) Bar plots showing the top 20 significantly enriched Gene Ontology (GO) Biological Process terms (p < 0.05), ranked by fold enrichment, for glial cluster 5 (g), cluster 6 (h) and cluster 19 (i), enriched in GB-transplanted hc-assembloids. Color intensity indicates the number of genes associated with each GO term (term size). Abbreviations: hc-human cerebral, ass-assembloid, tr-transplanted, logFC-log fold change, GO-Gene Ontology

We next asked whether these GB-enriched astroglial states shared a common transcriptional program. Differential gene expression analysis comparing each GB-enriched cluster with all remaining astroglial clusters revealed only 15 genes commonly upregulated across all five clusters (**Fig. 4e**; **Table S9**). A module score calculated from these shared genes was consistently enriched across all GB-associated astroglial clusters (**Fig. 4f**), defining a common pan-reactive astrocytic signature. Notably, this signature differed from previously described reactive astrocyte programs associated with traumatic brain injury^32^. Instead, it included genes previously implicated in glioblastoma biology, including *HMGA2-AS*, *SREBF2*, *USP34*, *CAB39* ^76–79^. Interestingly, several of the differentially expressed genes, including *SREBF2*, *CAB39*, *OTUD7B*, *USP34*, and *XRN1*, converge on pathways regulating metabolic adaptation, protein homeostasis, and RNA/protein turnover, indicating that metabolic reprogramming is a common feature of GB-induced reactive astrocytes and is part of a broader adaptive transcriptional response to the hc-assembloid niche. Consistent with this interpretation, Gene Ontology analysis of cluster 5, a cluster shared between two GB lines, revealed significant enrichment of metabolic processes (**Fig. 4g**). To determine whether this common reactive astrocyte signature is conserved in patient tissue, we analyzed its expression in a publicly available spatial transcriptomics dataset containing glioblastoma infiltration zones ^80^ (**Fig. S6h-m**). We first identified malignant regions using a malignancy score (**Fig. S6h,j,l**; **Table S8**) and subsequently assessed the expression of the reactive astrocyte signature derived from the 15 genes commonly upregulated in our hc-assembloid model. Notably, regions exhibiting a high reactive astrocyte signature strongly colocalized with areas of high malignancy score, corresponding to the tumor infiltration zone (**Fig. S6i,k,m**). These findings suggest that the reactive astrocyte signature identified in our model is conserved in patient tissue and support the notion that metabolic reprogramming of reactive astrocytes represents a common feature of the glioblastoma peritumoral microenvironment across distinct GB subtypes.

Although all GB-induced astroglial states shared this common transcriptional program, each GB subtype also induced distinct reactive signatures. GBM14NSC-enriched clusters (clusters 3 and 6) were characterized by genes associated with enhanced transcription, translation, and RNA processing (**Fig. 4h**; **Fig. S6d**) In contrast, LN229-enriched clusters (clusters 1 and 19) preferentially expressed genes involved in cell migration, extracellular matrix interactions, and immune responses (**Fig. 4i; Fig. S6e**), consistent with the highly invasive phenotype of MES1 like mesenchymal GB.

Taken together, these findings may indicate that distinct GB states induce reactive astrogliosis with subtype-specific characteristics.

To validate this hypothesis, we established primary glioblastoma cultures from surgical tumor explants representing distinct GB IDH-wildtype subtypes (**Fig. 5a**; **Table S7**). Cells were labelled with the lipophilic dye PKH26 (**Fig. 5a′, 5b-d**), allowing their visualization 3.5 weeks after transplantation (**Fig. 5e**). Following transplantation, primary GB cells formed compact tumor masses characterized by densely packed DAPI-positive nuclei surrounding a central region containing pyknotic nuclei, indicative of necrosis (**Fig. 5n, n′**). We first assess if the GB generated in the assembloids resemble the patient tumor tissue. Towards this end, human GB specimens were stained with hematoxylin and eosin to characterize growth patterns and tumor morphology (**Fig. 5f,j**). In addition, immunohistochemical staining for Olig2 was performed (**Fig. 5g,k**), with Olig2 serving as a surrogate marker of tumor cells in this context. Hc-assembloids transplanted with primary patient-derived tumor cells (NGLs) were analyzed in the same manner **(Fig. 5h,I,i,m)**. The NGL tumors within transplanted hc-assembloids recapitulated the diffuse growth pattern seen at the tumor border zones, whereas their centers exhibited pyknotic nuclei, apoptotic cells, and necrotic areas (**Fig. 5h,l**), consistent with a central deficit in nutrient and oxygen supply. We next evaluated the reactive gliosis in the PAs. Unlike LN229 and GBM14NSC tumors, NGL were infiltrated by hc-assembloid-derived microglia (**Fig. 5n, n′**). Importantly, the tumor bulk was surrounded by sparsely distributed PKH26-positive invading tumor cells within regions displaying pronounced GFAP upregulation, consistent with the establishment of a PA and the induction of reactive gliosis (**Fig. 5n′′**). Remarkably, the three primary GBs induced distinct patterns of astrocyte activation and microglial accumulation in the PA (**Fig. 5o-q**). These subtype-specific glial responses closely mirrored the pattern of gliosis observed in the corresponding patient specimens (**Fig. 5r-t**), with strong correlations between the hc-assembloid model and patient tissue (R^2^ = 0.661 for reactive astrocytes and R^2^ = 0.920 for microglia)**(Fig. 5u,v).**

**Figure 5.**
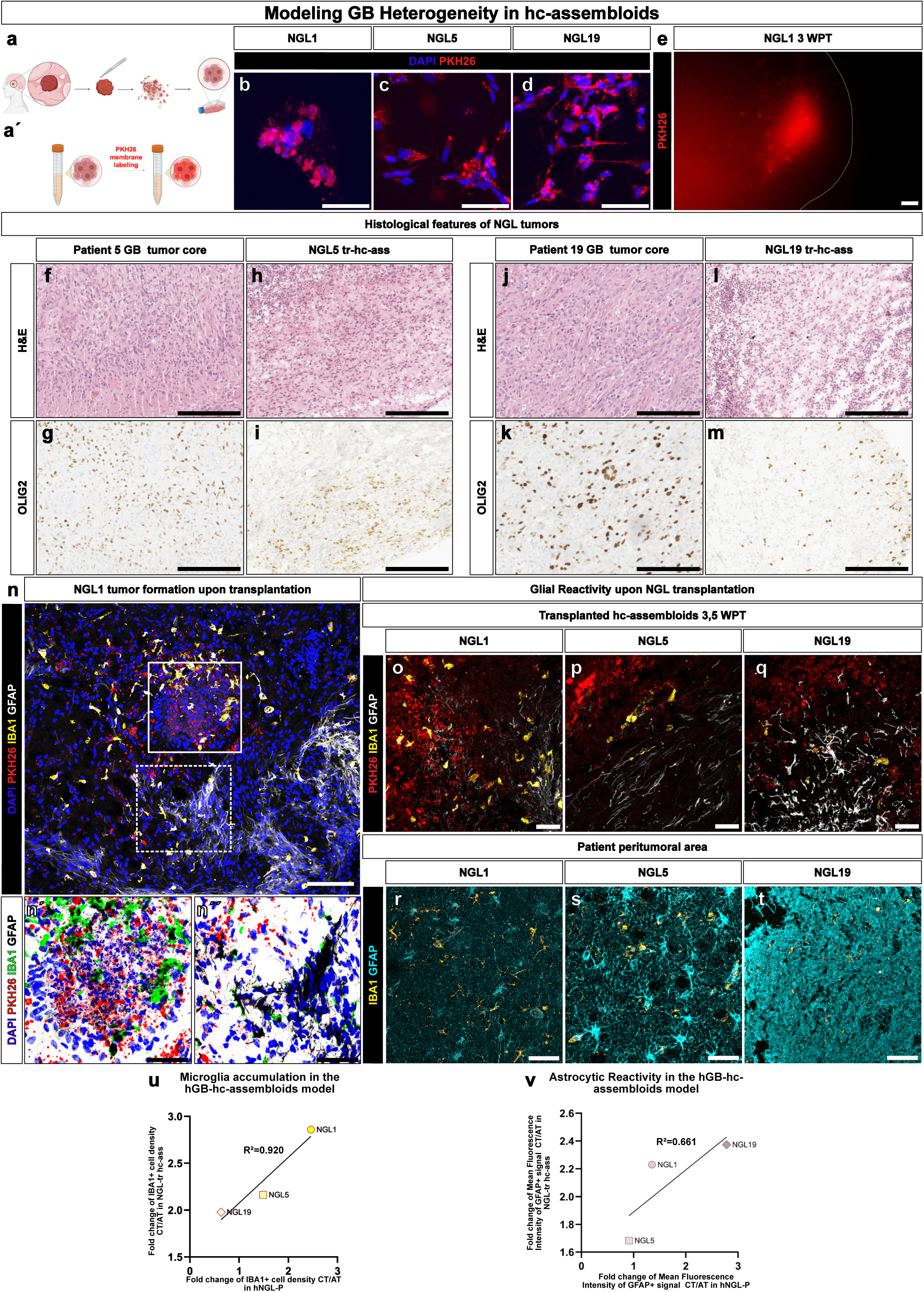
Establishment of the peritumoral area in hc-assembloids following transplantation of patient-derived GB cells. (a, á) Scheme depicting experimental workflow for derivation (a) and labelling (á) of primary, patient derived GB cells (NGL). (b-d) Micrographs depicting NGL-PKH26 labelled, plated cells prior to transplantation for different GB cells, NGL1 (b), NGL5 (c) and NGL19 (d). (e) Fluorescence micrograph showing NGL1 PKH26-labelled cells within hc-assembloid three weeks post transplantation (3 WPT). (f-m) Histological features of NGL tumors. (f, g, j, k) Micrographs depicting the histology of the parental patient tumors from which the primary lines NGL5 (f, g) and NGL19 (j, k) were derived. Formalin-fixed, paraffin-embedded (FFPE) sections of the GB tumor core of patient 5 (f, g) and patient 19 (j, k) were stained with hematoxylin and eosin (H&E) (f, j) and for OLIG2 (g, k), which serves as a surrogate marker of tumor cells. (h, i, l, m) Micrographs depicting FFPE sections of NGL5-(h, i) and NGL19-(l, m) transplanted hc-assembloids, processed and stained in parallel with H&E (h, l) and for OLIG2 (i, m). (n) Micrograph depicting NGL1 tumor formation 3.5 weeks after transplantation in hc-assembloids. Panels (ń, ń′) present magnified images of NGL1 tumor bulk (ń, continuous white box) and PA (ń′, dashed white box). (o-q) Micrographs demonstrating host glial reactivity within transplanted hc-assembloids 3,5 weeks post-transplantation for NGL1 (o), NGL5 (p), and NGL19 (q) cells. (r-t) Representative micrographs showing glial reactivity in the PA of the corresponding patient tissues from which NGL1 (r), NGL5 (s), and NGL19 (t) were derived. (u, v) Dot plots showing the correlation between microglial enrichment (u) and enrichment of GFAP signal (v) in the PA of hc-assembloids and the corresponding patient tissue. Enrichment was calculated relative to tissue regions distant from the peritumoral area. Individual data points represent the mean of 3-4 transplanted hc-assembloids derived from two independent differentiation batches for the hc-assembloid model, and the mean of two histological sections per patient for the corresponding human tissue. Normality was assessed using the Shapiro–Wilk test. Scale bars: 200 μm in f-m, 50 μm in b-d, ń, ń′, o-t and 100 μm in c, d. Abbreviations: NGL-Ninkovic Lab Glioblastoma Line, hc-human cerebral, H&E-hematoxylin and eosin, tr-transplanted, WPT-weeks post transplantation

Collectively, these findings demonstrate that the hc-assembloid model faithfully recapitulates GB patient-specific reactive gliosis and reproduces the glial responses observed in patient peritumoral tissue, providing a physiologically relevant platform to investigate subtype-dependent tumor–brain interactions.

#### GB-Transplanted hc-Assembloids as a Discovery Platform

A defining feature of glioblastoma progression is the extensive remodelling of cell–cell communication within both the tumor bulk and the surrounding PA ^81^. To identify potential intercellular signalling networks in our model, we applied the CellPhoneDB pipeline ^70^, which infers putative ligand–receptor interactions based on cell-type composition and the expression of known ligand–receptor pairs in the scRNA dataset. We examined putative ligand–receptor interactions between GB cells and astrocytes. Comparison of LN229- and GBM14NSC-transplanted hc-assembloids with media-transplanted controls revealed extensive remodelling of intercellular communication (**Fig. 6a**). The majority of newly predicted interactions involved extracellular matrix (ECM) components and their cognate receptors (**Fig. 6b**), consistent with the concept that reciprocal interactions between GB cells and the surrounding niche shape both tumor progression and glial reactivity. Notably, only a subset of astrocyte clusters, particularly clusters 5, 10, and 19, engaged in a disproportionately large number of predicted interactions, suggesting that these populations function as signalling hubs within the peritumoral microenvironment (**Fig. 6a**).

**Figure 6.**
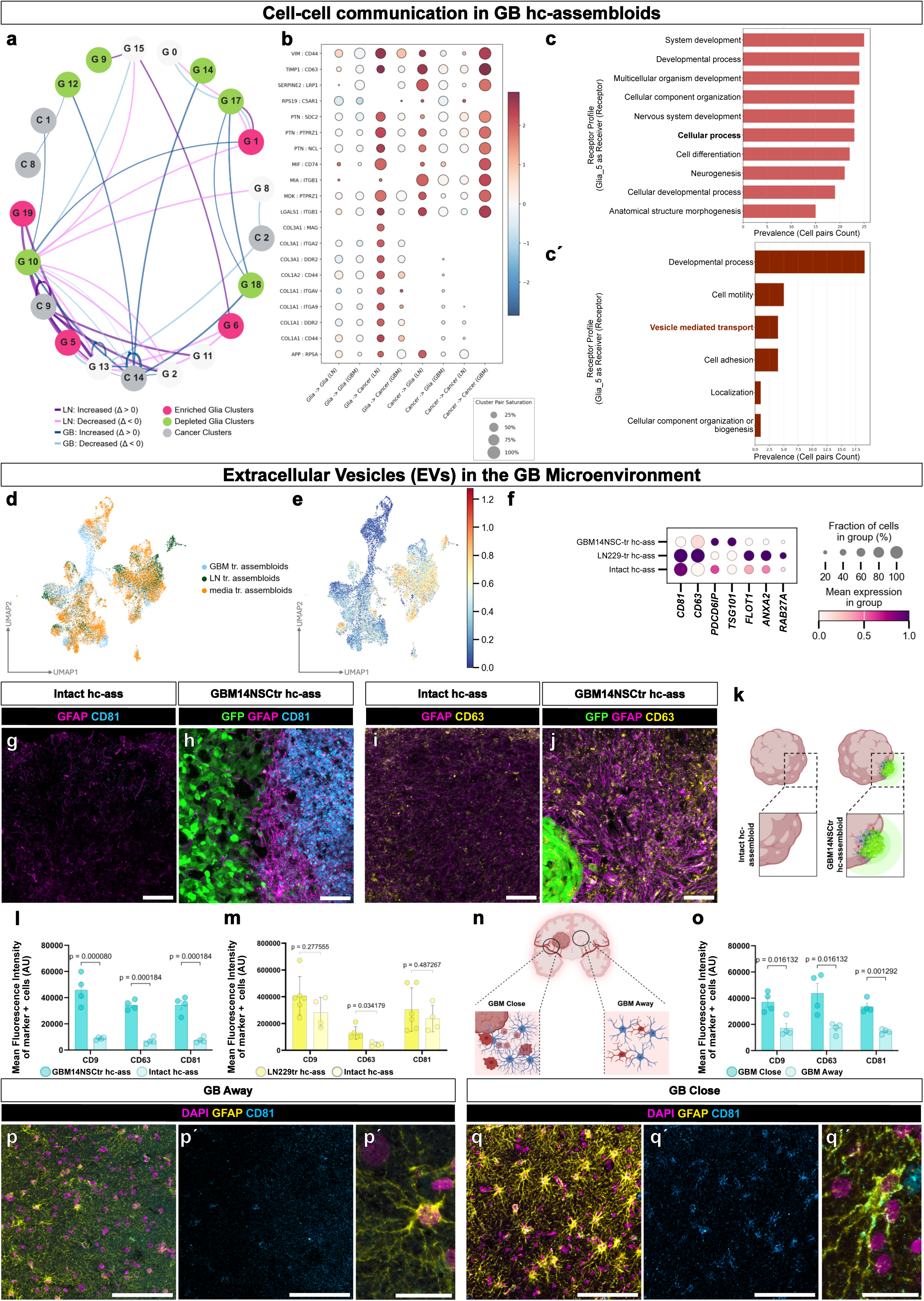
Community-based analysis of putative interactions of GB and astroglial cells. (a) Sankey plot depicting communication analysis among glial and cancer cell clusters. Nodes represent glial and cancer clusters and edge thickness indicates the number of predicted ligand-receptor pairs between clusters, with thicker edges indicating a greater extent of potential cell-cell communication. Edge color denotes the direction of pathway regulation in LN229 transplanted (purple, increased; pink, decreased) and GBM14NSC transplanted (dark blue, increased; light blue, decreased) hc-assembloids. (b) Dot plot depicting the top 20 ligand and receptor pairs between glia and cancer clusters from the network in panel a. (c) GO classification of ligand–receptor interactions received by the glia cluster 5. (c′) Hierarchical decomposition of the *cellular process* GO category identified in (c), highlighting the major biological processes represented within the cellular process term. (d) UMAP plot depicting astroglial cells colored by condition. (e) UMAP plot presenting the distribution of EV gene expression score across astroglial cells. (f) Dot plot showing the expression of selected, EV-associated genes comprising the EV expression score (e), in GBM14NSC-, LN229-, and intact hc-assembloids. (g-j) Micrographs depicting GFAP-positive astrocytes and expression of the EV markers CD81 (g, h) and CD63 (i, j) in intact assembloids (h, j) and in the PA of transplanted (h, j) hc-assembloids. (k) Scheme illustrating hc-assembloid regions used for EV quantifications in (l) and (m). (l, m) Dot plots showing quantification of signal intensity of the EV markers CD9, CD63, and CD81 in the PA of GBM14NSC-transplanted (l) and LN229-transplanted (m) hc-assembloids relative to intact hc-assembloids. Data are presented as mean ± Standard deviation (SD). Individual dots represent hc-assembloids obtained from three (l) and four (m) independent batches. Normality was assessed using the Shapiro-Wilk test and statistical analysis was performed using multiple unpaired t-tests. (n) Schematic illustrating the position of the human tissue relative to the tumor used for analysis in panels (o, p-p′, q-q′). (o) Dot plot illustrating the signal intensity of EV markers CD9, CD63, and CD81, in areas proximal and distal to the tumor. Data are presented as mean ± Standard deviation (SD). Each dot represents a different patient. Normality was assessed using the Shapiro-Wilk test and statistical analysis was performed using multiple unpaired t-tests. (p-q′′) Micrographs showing GFAP-positive astrocytes and the expression of the EV marker CD81 distant from the tumor (p, p′′) and near the tumor (q, q′′). (p′′) and (q′′) are magnified views of single CD81+ astrocytes in p and q, respectively. Scale bars: 100 μm in p, p′, q, q′ and 50 μm in g-j′, p′′and q′′. Abbreviations: EVs-Extracellular Vesicles, GB-Glioblastoma, hc-human cerebral, ass-assembloid, tr-transplanted

Because astrocyte cluster 5 was strongly enriched in both LN229- and GBM14NSC-transplanted hc-assembloids, we investigated the biological processes associated with its predicted ligand-receptor interactions. Gene Ontology analysis revealed enrichment of pathways involved in nervous system development, neurogenesis, cell differentiation, and vesicle-mediated transport (**Fig. 6c, c′**). These findings suggested that reactive astrocytes may not only respond to extracellular vesicles (EVs) released by GB cells, as previously reported ^82^,but may themselves actively participate in bidirectional communication within the PA through EV secretion. To test this hypothesis, we compared the expression of EV-associated genes in reactive astrocytes from GB-transplanted and astrocytes from media-transplanted hc-assembloids in the scRNAseq dataset. Reactive astrocytes from GB-transplanted hc-assembloids exhibited significantly increased expression of EV-related genes, reflected by a marked enrichment of the corresponding gene expression signature (**Fig. 6d-f**).

We next validated these findings at the protein level by immunohistochemistry for the EV markers CD81, CD63, and CD9 (**Fig. 6g-m**). All three markers were significantly enriched in the PA of GB-transplanted hc-assembloids compared with intact tissue following transplantation of either GBM14NSC or LN229 cells (**Fig. 6l,m**). Importantly, EV marker expression strongly overlapped with GFAP-positive reactive astrocytes (**Fig. 6h,j**), suggesting that reactive astrocytes are an important source of extracellular vesicles within the PA. To our knowledge, increased EV production by reactive astrocytes in the PA has not previously been demonstrated in patient tissue. We therefore examined the expression of CD81, CD63, and CD9 in human glioblastoma samples by comparing tissue immediately adjacent to the tumor with more distant brain regions from the same patient (**Fig. 6n**). Remarkably, the PA displayed significantly higher expression of all three EV markers (**Fig. 6o-q′′**). Similar to the hc-assembloid model, EV marker expression strongly colocalized with GFAP-positive astrocytes (**Fig. 6q-q′′**), demonstrating that reactive astrocytes in patient tissue likewise exhibit increased EV production.

To determine whether EV secretion contributes functionally to the maintenance of the PA, we pharmacologically inhibited EV release using the ROCK inhibitor Y-27632, which has been reported to suppress EV secretion ^83^ (**Fig. 7a**). Nanoparticle tracking analysis (NTA) demonstrated that one week of Y-27632 treatment almost completely abolished EV release, whereas vehicle-treated LN229- and GBM14NSC-transplanted hc-assembloids released vesicles with an average diameter of approximately 200 nm (**Fig. 7b,b′**). Transmission electron microscopy confirmed that these particles displayed the characteristic bilayer membrane morphology of extracellular vesicles ^84^ including exosomes and microvesicles (**Fig. 7c,c′**).

**Figure 7.**
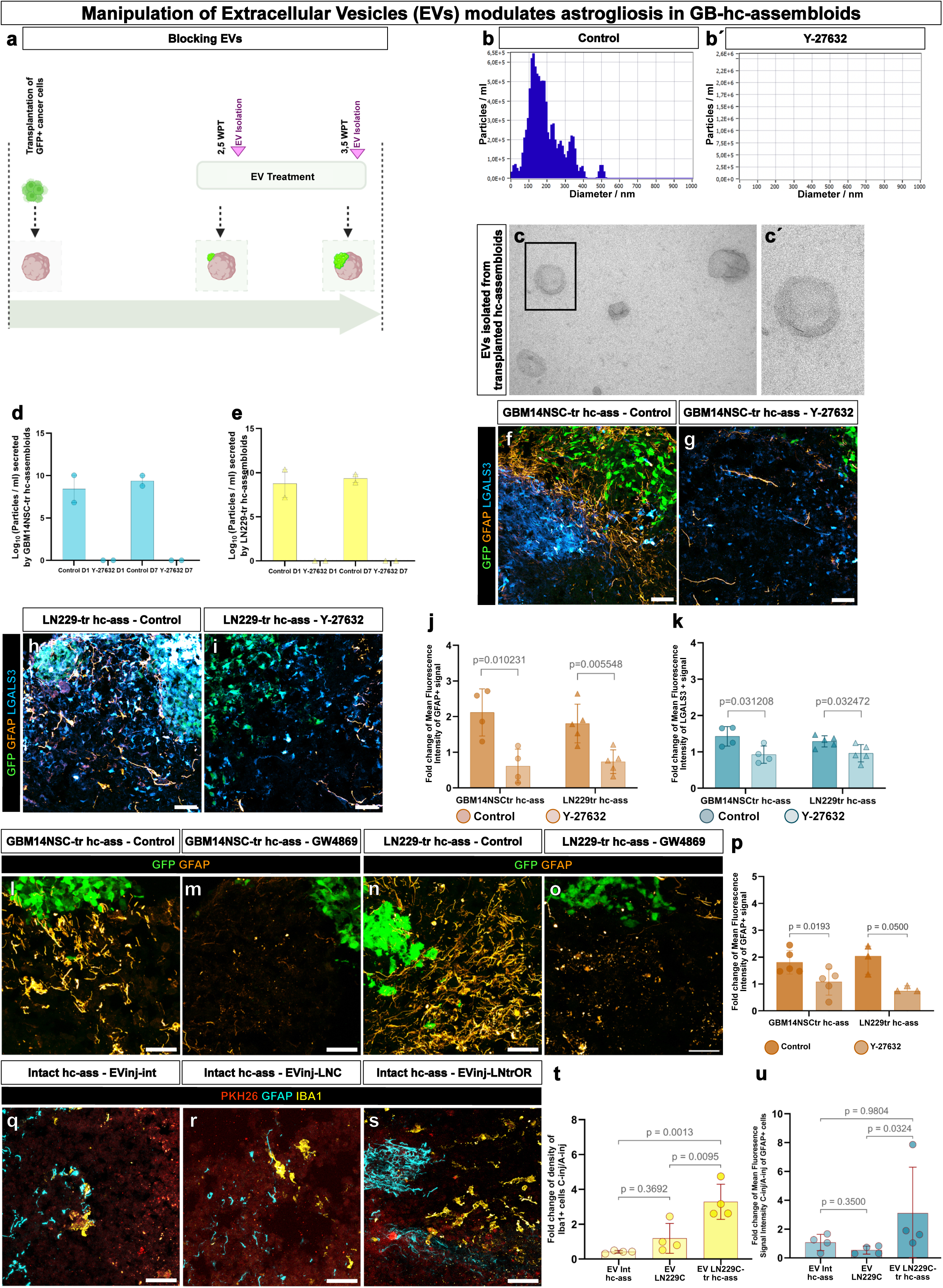
The role of Extracellular Vesicles (EV) in establishing reactive astrogliosis in the peritumoral area. (a) Schematic workflow for blocking EVs in transplanted hc-assembloids. (b-b′) Nanoparticle Tracking Analysis (NTA) curves showing particle concentration as function of particle diameter for particles isolated from GBM14NSC-transplanted hc-assembloids under control (b) and Y-27632-treated (b′) conditions. (c, c′) Micrographs depicting transmission electron microscopy (TEM) image of EVs isolated from transplanted hc-assembloid culture media. Panel (c′) is a magnified image of the black boxed area in (c). (d, e) Bar plots showing the concentration of EVs secreted by GBM14NSC-(d) and LN229-transplanted hc-assembloids (e) following Y-27632 and corresponding vehicle treatment after 1 and 7 days of treatment. Each dot represents an independent hc-assembloid differentiation batch, with EVs collected from the culture medium of 2-3 hc-assembloids. (f-i) Micrographs depicting GFAP and LGALS3 astroglial reactivity in the peritumoral area of GBM14NSC-(f, g) and LN229-(h, i) transplanted hc-assembloids after vehicle (control, f, h) and Y-27632 inhibitor treatment (g, i). (j, k) Dot plots presenting the intensity of GFAP-(j) and LGALS3-(k) signal following Y-27632 and control treatment, calculated as the ratio of signal close to the tumor and signal distant from the tumor. Data are presented as mean ± Standard deviation (SD) and each dot represents individual hc-assembloid. Hc-assembloids were obtained from 2 independent batches for all conditions. Normality was assessed using the Shapiro-Wilk test and statistical analysis was performed using multiple unpaired t-tests. (l-o) Micrographs depicting GFAP astroglial reactivity in the peritumoral area of GBM14NSC-(l, m) and LN229-(n, o) transplanted hc-assembloids treated with vehicle (control, l, n) and GW4869 inhibitor (m, o). (p) Dot plot showing the GFAP signal intensity, expressed as the ratio between the PA and area distant from the tumor, following treatment with GW4869 or the corresponding vehicle control. Data are presented as mean ± Standard deviation (SD) and each dot represents an individual hc-assembloid, obtained from 3 independent differentiations. Normality was assessed using the Shapiro-Wilk test and statistical analysis was performed using multiple unpaired t-tests. (q-u) Reactive gliosis 5 days upon injection of intact hc-assembloids with EVs derived from intact hc-assembloids (q), LN229 cancer cells (q) and LN229 transplanted hc-assembloids (s). (t) Dot plot showing the GFAP signal intensity, expressed as the ratio between the region close to injection and region distant from the injection site in EV-injected hc-assembloids. (u) Dot plot depicting the density of IBA-positive cells, expressed as the ratio between the region close to injection and region distant from the injection site in injected hc-assembloids. Data are presented as mean ± Standard deviation (SD) and each dot represents an individual hc-assembloid. Normality was assessed using the Shapiro-Wilk test and statistical analysis was performed using Kruskal-Wallis (t) and One-way Anova (u) accordingly. Scale bars: 200 μm in c, c′ and 50 μm in f-I and l-o. Abbreviations: EVs-Extracellular Vesicles, GB-Glioblastoma, tr-transplanted, hc-human cerebral, ass-assembloid, WPT-Weeks Post Transplantation, D1-Day1, D7-Day7.

To investigate the consequences of EV inhibition on reactive gliosis, hc-assembloids were first transplanted with LN229 or GBM14NSC cells and cultured for 2.5 weeks to allow establishment of the PA. Cultures were then treated with either Y-27632 or vehicle for an additional seven days before assessing astrocyte activation by GFAP immunostaining and extracellular matrix remodelling by LGALS3 immunostaining (**Fig. 7a,f-i**). Inhibition of EV secretion (**Fig. 7d,e**) resulted in a significant reduction of both GFAP and LGALS3 expression compared with vehicle-treated controls (**Fig. 7j-k**). Similar results were obtained using GW4869 (**Fig. 7l-p**), a neutral sphingomyelinase inhibitor that blocks ceramide-dependent EV biogenesis ^83,85,86^, confirming that these effects were attributable to inhibition of EV production rather than off-target effects of ROCK inhibition. We next tested whether PA-derived EVs were sufficient to induce glial reactivity in intact hc-assembloids (**Fig. 7q–u**). EVs derived from intact hc-assembloids, LN229 cultures, or LN229-transplanted hc-assembloids were labelled with a membrane dye and subsequently microinjected into intact hc-assembloids at culture day 129. Five days later, we assessed the local glial response around the injection site, identified by the presence of the lipophilic dye (**Fig. 7q–s**). Injection of EVs derived from LN229-transplanted hc-assembloids induced increased accumulation of IBA1-positive microglia compared with EVs derived from intact hc-assembloids or LN229 cells alone (**Fig. 7t**). In addition, we observed a non-significant trend toward increased GFAP signal around the injection site following injection of EVs derived from LN229-transplanted hc-assembloids compared with EVs from intact hc-assembloids or LN229 cells alone (**Fig. 7u**).

Collectively, these findings demonstrate that extracellular vesicles are produced not only by GB cells but also by reactive astrocytes within the PA, where they contribute to multidirectional intercellular communication. Moreover, pharmacological inhibition of EV secretion disrupts both reactive astrogliosis and extracellular matrix remodelling, indicating that EV signalling is essential for maintenance of the peritumoral microenvironment. Together, these results highlight the hc-assembloid model as a powerful discovery platform capable of identifying previously unrecognized features of the glioblastoma PA that can subsequently be validated in patient tissue.

#### Modularity of the hc-Assembloid System for Deciphering the GB Niche

GB progression is driven by complex and often redundant interactions between multiple cellular compartments within the tumor microenvironment ^19,24^. Our hc-assembloid model provides a modular platform that allows for the controlled integration of different cell types involved in GB progression. This adaptability enables a systematic dissection of functional roles, offering critical insights into the individual and collective contributions of various cellular players.

To demonstrate this modularity, we incorporated iPSC-derived monocytes (i-monocytes) into the hc-assembloid system. Monocytes and microglia share overlapping roles in tumor-associated immune responses, yet their precise contributions remain difficult to delineate in vivo due to functional redundancy and lack of distinct markers ^87–89^. We first generated i-monocytes using a modified Cao et al. protocol ^90^ from three different iPSC lines (**Fig. S7a**). Generated i-monocytes express comparable levels of CD14 and CD16 independently of the iPSC line (**Fig. S7b**), with a purity of approximately 40% (**Fig. S7b**). Notably, the i-monocytes expressing CD14 and IBA1 (**Fig. S7c**-**d′′**) demonstrated functional maturation by actively phagocytosing pHrodo™ BioParticles™ Conjugates (**Fig. S7e**-**f′′**).

To assess whether i-monocytes could integrate into hc-assembloids, we inoculated hc-assembloids with RFP-expressing iPSC-derived i-monocytes and tracked the infiltration of i-monocytes into hc-assembloids (**Fig. S7g**). RFP-positive cells were observed within hc-assembloids, confirming their successful integration in both intact and tumor bearing hc-assembloids (**Fig. 7h, 7i, Video S5, S6**). Importantly, all RFP-positive cells infiltrating hc-assembloids expressed CD14 and MPEG1, confirming their monocyte identity (**Fig. S7h′-h′′′, i’-i’’’**). When co-cultured with GB14NSC-transplanted hc-assembloids, i-monocytes preferentially populated GB-containing regions (**Fig. S7i**). In contrast to microglia, which accumulate primarily in the PA (**Fig. S5f-f′′**), i-monocytes were found in both the tumor core and PA (**Fig. S7i, Video S6**), mirroring patterns observed in patient samples ^91,92^. These findings highlight the hc-assembloid system’s ability to model immune-tumor interactions in a controlled setting, allowing for the study of immune and stromal components in the GB PA. Beyond monocytes, this modularity can be extended to incorporate endothelial cells and other peripheral immune populations, creating a highly adaptable and physiologically relevant platform for investigating tumor-stroma interactions.

The modular approach enables targeted manipulation of the GB microenvironment, facilitating the identification of key cellular players and molecular pathways involved in disease progression. Ultimately, this system serves as a powerful tool for studying GB biology and testing novel therapeutic strategies aimed at disrupting tumor-supporting interactions.

## Discussion

Glioblastoma (GB) is characterized by a profoundly complex tumor microenvironment that extends far beyond the heterogeneous tumor mass and includes an intricate peritumoral area (PA) ^24,93^. The PA comprises a diverse collection of cell types, including infiltrative immune cells (e.g., T-cells, B-cells, dendritic cells, monocytes, macrophages, natural killer cells, and neutrophils), various glial cells (astrocytes, microglia, and oligodendrocytes), neurons, and cancer stem cells ^94–96^. This cellular diversity fosters a highly dynamic microenvironment marked by complex cell–cell interactions, intricate immune modulation, and reactive gliosis, all of which contribute to the aggressive phenotype and therapeutic resistance observed in GB. Given these complexities, pinpointing the individual contributions of specific cellular populations is challenging. Therefore, developing a human in vitro model that faithfully recapitulates this multifaceted microenvironment is essential for advancing our understanding of GB biology. Despite substantial advances in organoid-based GB modelling, existing systems remain limited in their ability to simultaneously reproduce the cellular complexity of the human brain microenvironment and the spatially defined interaction of patient-derived GB with initially non-neoplastic brain tissue. Genetically engineered organoid models provide a human three-dimensional tissue context and enable controlled analysis of defined oncogenic alterations, but tumor formation occurs intrinsically within the organoid and is driven by a restricted set of experimentally introduced mutations ^97–99^. Consequently, although these models are particularly valuable for dissecting the contribution of individual driver alterations to tumor initiation and progression, they do not reproduce the full somatic mutational complexity, plasticity, transcriptional heterogeneity, and epigenetic landscape of patient-derived GB. Conversely, cerebral organoid co-culture models preserve patient-derived tumor characteristics and permit direct analysis of GB invasion into human neural tissue ^97,100^, but typically introduce relatively large tumor-cell populations onto the organoid surface, resulting in variable and spatially dispersed tumor growth and limiting the availability of clearly defined tumor-free, peritumoral, and tumor-containing regions within the same tissue. More advanced microfluidic systems can incorporate vascular compartments, extracellular-matrix components, and blood-brain barrier-like properties ^101^, but increasing these individual aspects of physiological complexity does not necessarily recreate the full cellular composition and organization of the human brain parenchyma. Similarly, immune-competent approaches such as iHOTT partially address the absence of immune components by incorporating autologous peripheral blood mononuclear cells into cortical organoid–GB co-cultures; however, the use of circulating immune cells rather than a resident microglial compartment, the relative underrepresentation of myeloid cells, and the short experimental window limit representation of the myeloid-rich and dynamically evolving GB microenvironment ^102^. This limitation is particularly relevant because brain-resident microglia represent a distinct component of the tumor microenvironment that can emerge within appropriately generated and sufficiently matured cerebral organoid systems ^28^. Thus, although individual platforms reproduce important aspects of GB initiation, invasion, immune interactions, vascular barriers, or therapeutic responses, they generally capture these features in isolation rather than within a single model that combines a complex human neural, glial, and resident immune microenvironment with spatially controlled interactions between patient-derived GB and surrounding non-neoplastic tissue. This limitation is particularly important for studying the PA, where reciprocal interactions between infiltrating tumor cells and resident neural, glial, and immune populations are likely to shape both tumor behavior and the response of the surrounding brain tissue. In order to model these interactions, we employed human cerebral assembloids (hc-assembloids) as a platform to model the GB microenvironment. Our approach was designed to replicate key cellular populations and structural elements of the human brain, including neuronal progenitors, neurons, astrocytes, microglia, and oligodendrocyte lineage cells. This inherent cellular heterogeneity renders the hc-assembloid model particularly well suited for GB transplantation studies, as it facilitates the formation of anatomically distinct regions within the hc-assembloid (a tumor core, a peritumoral zone, and tumor-free areas) that closely mimic the patient tumor landscape ^5^. A significant advantage of our transplantation paradigm is that it promotes the formation of localized tumor niches with infiltrating front within the hc-assembloids, thereby providing spatial context that is often lacking in traditional co-culture methods ^103^. Our approach also facilitates interactions with the non-tumor parenchymal cells that constitute the PA – a unique concept so far to our knowledge ^99,100,104,105^. Although our hc-assembloid system may contain immature cells, the maturity of astrocytes and microglia appears sufficient to induce typical niche reactions, including upregulation of GFAP and morphological changes in microglia observed in patients ^106^ that we could also validate using primary patient tissue. Moreover, the microglia in our system express “sensome genes” that have previously been observed only in vivo or after the transplantation of human organoids into an in vivo context ^39^ and is essential for the PA formation further substantiating the versatility of our model. Importantly, our model system, even in its simplest version, allows us not only to study the interaction of parenchymal glia with GB but also to explore the interplay with tumor-associated interneuron populations, as evidenced by the presence of GB-relevant calbindin-positive interneurons ^107^. We start by transplanting 2,000 GB cells. Over time, these cells grow from small tumor foci into a large mass that interacts with the surrounding tissue, closely mimicking what happens in patients. GB cells in the hc-assembloids exhibit more pronounced tumor subtype characteristics compared to traditional GB cell cultures, consistent with in vivo observations and highlighting the plasticity of GB cells under microenvironmental influences ^45,108^. The microenvironment is defined not only by its cellular composition but also by its biophysical properties. Tissue stiffness, for instance, has been shown to influence tumor cell behavior by promoting invasion and resistance to therapy ^23,109^. In our study, we measured the mechanical properties of tumor-bearing hc-assembloids and found that the tumor core exhibits a stiffness (Young’s modulus) of 1.6 kPa, in good agreement with patient-derived tumor measurements ^61^. Concomitantly, we observed upregulation of activation markers in astrocytic and microglial populations within tumor-bearing organoids, specifically, GFAP, CD44, and LGALS3 in astrocytes. These markers indicate a robust reactive gliosis response that parallels the GB-induced neuroinflammatory processes observed in patients ^20^. Notably, control experiments using hiPSC-derived neural progenitor cells (NPCs) in place of malignant cells failed to induce a similar reactive response, confirming that the observed gliosis is tumor-specific.

Another novel aspect of our study is the elevated expression of extracellular vesicle (EV) markers in peritumoral astrocytes. EVs have recently emerged as pivotal mediators of intercellular communication within tumors, capable of transferring oncogenic signals and facilitating tumor progression ^82,110,111^. Our findings suggest that reactive glial cells may employ EVs to communicate either with other parenchymal cells in the PA or/ and directly with GB cells, positioning EV-mediated signalling as a promising therapeutic target. Another important feature of our system is its modularity, as demonstrated by the successful incorporation of iPSC-derived monocytes into the hc-assembloid culture. These i-monocytes rapidly infiltrate the hc-assembloid tissue, underscoring the model’s potential for studying specific immune-tumor interactions that shape the PA environment ^89,112^. Future iterations of the model could integrate additional immune cell types, such as GB-relevant T-cells or dendritic cells ^113,114^, thereby providing a comprehensive platform to study immune dynamics in GB.

Despite its many strengths, our model is not without limitations. The most notable is the absence of vascularization which is a critical feature of the in vivo GB microenvironment ^115^. Angiogenesis is fundamental for tumor growth as it supplies nutrients and facilitates tumor cell migration ^116^. Although our current model lacks endothelial cells and hypoxic gradients, recent advances in vascularized organoid technology ^117^ hold promise for addressing this limitation. Another approach to overcome the limitation of the well-known hypoxic core in hc-assembloids ^118^, would be to combine our model with already reported slicing organoid protocols ^119,120^. Additionally, while our hc-assembloids exhibit a diverse array of glial cells, questions remain regarding their functional maturity. Enhancing differentiation protocols to promote further glial maturation or incorporating adult-derived glial cells may yield a more accurate simulation of the mature brain environment and enhance the model’s translational relevance.

In summary, our hc-assembloid model represents a promising and innovative platform for studying GB within a complex, brain-like microenvironment. By faithfully replicating key features of the tumor milieu—including cellular diversity, mechanical properties, immune responses, and intercellular communication—this model offers substantial advantages for both fundamental and translational GB research. Future refinements, such as the integration of vascular networks and enhanced glial maturation, will further elevate the model’s utility, paving the way for novel therapeutic discoveries and personalized medicine approaches in neuro-oncology.

## STAR METHODS

Detailed methods are provided in the online version of this manuscript and include the following:

### RESOURCE AVAILABILITY

#### Lead contact

Requests for further information and resources should be directed to and will be fulfilled by the lead contacts: S. Kalpazidou, P. Moysidou-Tsiorva, Anna Danese and Jovica Ninkovic

#### Materials availability

The HMGU1 and HMGU12 iPSC lines, generated from human foreskin fibroblasts at the iPSC Core Facility, Helmholtz Center Munich, are available upon request from the iPSC Core Facility, Helmholtz Center Munich. The HMGU1 iPSC line, which expresses Red Fluorescent Protein (RFP), was provided by Prof. Dr. Silvia Cappello from the Department of Physiological Genomics at the Biomedical Center of LMU. The GM25256 (WTC-11) iPSC line was purchased by Coriell Institute for Medical Research.

Human glioblastoma (GB) cell lines used in this study: The GBM14NSC cell line, derived from a human glioblastoma biopsy, is available upon request from Prof. Dr. Rainer Glaß at LMU University Hospital, Neurosurgical Research. The LN-229 (CRL-2611) cell line was kindly provided by Prof. Dr. Jürgen Schlegel from the Neuropathology Department at the Technical University of Munich (TUM).

#### Materials And Equipment

| Low FGF Medium |  |  |
| --- | --- | --- |
| Reagent | Final Concentration | Amount |
| DMEM/F12 | N/A | 40 mL |
| KOSR | - | 10 mL |
| ES-quality FBS | 3% | 1.5 mL |
| GlutaMAX | 1X | 0.5 mL |
| MEM-NEAA | 1X | 0.5 mL |
| 50 mM 2-Mercaptoethanol solution | 0.1 mM | 100 µL |
| **Total** | N/A | 50 mL |
| This medium can be stored at 4°C for up to 2 weeks. |  |  |

Note: Just before use, add basic fibroblast growth factor (bFGF) to the medium at a final concentration of 4 ng/ml and ROCK Inhibitor Y27632 (RI) at a final concentration of 0.05 mM.

| Neuronal Induction Medium |  |  |
| --- | --- | --- |
| Reagent | Final Concentration | Amount |
| DMEM/F12 | N/A | 100 mL |
| N2 supplement | 1X | 1 mL |
| GlutaMAX | 1X | 1 mL |
| MEM-NEAA | 1X | 1 mL |
| Heparin | 0.1 µg/mL | 10 µL |
| **Total** | N/A | 100 mL |
| This medium can be stored at 4°C for up to 2 weeks. |  |  |

| Improved Cerebral Organoid Differentiation Medium without Vitamin A (IDM-A) |  |  |
| --- | --- | --- |
| Reagent | Final Concentration | Amount |
| DMEM/F12 | - | 125 mL |
| Neurobasal | - | 125 mL |
| N2 supplement | 0.5X | 1.25 mL |
| B27 supplement (without Vitamin A) | 1X | 5 mL |
| Insulin | 2.5 µg/ml | 62.5 µL |
| 50 mM 2-Mercaptoethanol solution | 0.05 mM | 250 µL |
| GlutaMAX | 1X | 2.5 mL |
| MEM-NEAA | 0.5X | 1.25 mL |
| P/S (Penicillin/Streptomycin) | 1X | 2.5 mL |
| **Total** | N/A | 250 mL |
| This medium can be stored at 4°C for up to 2 weeks. |  |  |

| Improved Cerebral Organoid Differentiation Medium with Vitamin A (IDM+A) |
| --- |

| Reagent | Final Concentration | Amount |
| --- | --- | --- |
| DMEM/F12 | - | 125 mL |
| Neurobasal | - | 125 mL |
| N2 supplement | 0.5X | 1.25 mL |
| B27 supplement | 1X | 5 mL |
| Insulin | 2.5 µg/ml | 62.5 µL |
| 50 mM 2-Mercaptoethanol solution | 0.05 mM | 250 µL |
| GlutaMAX | 1X | 2.5 mL |
| MEM-NEAA | 0.5X | 1.25 mL |
| P/S (Penicillin/Streptomycin) | 1X | 2.5 mL |
| **Total** | N/A | 250 mL |
| This medium can be stored at 4°C for up to 2 weeks. |  |  |

#### Data and code availability

Microscopy data and reported Single-cell RNA-seq in this paper will be shared by the lead contacts upon request.

This paper utilizes published spatial data generated by Greenwald et al. and publicly available on Zenodo ^80^.

All original code has been deposited at the Ninkovic Lab Github repository (available for reviewers upon request; will be public after manuscript has been accepted for publication). Any additional information needed to reanalyse the data in this paper can be obtained from the lead contacts upon request.

### EXPERIMENTAL MODELS AND STUDY PARTICIPANT DETAILS

#### iPSC Culture

Male iPSCs were used in the study as described in the Key Resources Table. Feeder-free iPSCs were maintained at 37°C, 5% CO_2_ and ambient oxygen levels on Geltrex coated wells of 6-well plates (Sardtedt, cat. no. 83.3920) with daily changes of the medium with fresh mTeSR. To remove spontaneously differentiating iPSCs, the iPSC colonies were assessed daily using the EVOs microscope. Differentiated parts of colonies were gently scratched using a 100 ul sterile pipette tip to avoid disturbing the remaining colonies. iPSCs colonies were grown until they reached 70–80 % confluency. Prior to splitting or thawing iPSCs, 6-well plates were coated with Geltrex (1:100 dilution in cold mTeSR) to provide a surface for the attachment of iPSCs. Plates were coated with Geltrex for at least 45 minutes at 37°C with 5% CO2. For passaging, iPSCs colonies were treated with 0.1% (vol/vol) collagenase IV (in PBS 1X) for 6-7 minutes at 37°C with 5% CO_2_ and then collagenase was replaced with fresh mTeSR. Using a 100 μl pippete tip colonies were scratched into smaller pieces and then scraped gently using a cell lifter. Detached colonies were gently distributed using a 5 ml serological pipettes into new, coated 6-well plates with fresh prewarmed mTeSR.

#### Generation of iPSC derived Neural Progenitor Cells (i-NPCs)

Neural progenitor cells (NPCs) were generated following an adapted protocol from Zink et al. (2021)^68^, utilizing the HMGU1-RFP iPSC line. Upon reaching 80% confluency, iPSC colonies were washed with 1X PBS and incubated with 1 mL of Accutase for 4 – 5 minutes at 37°C to achieve single-cell dissociation. Following incubation, 5 mL of 1X PBS were added, cells were collected in a conical tube and centrifuged at 120 × g for 5 minutes. The supernatant was removed, cells were resuspended in 500 μL of low-FGF medium, counted, and seeded at a density of 9 × 10³ cells per well in a low-attachment 96-well U-bottom plate using a multichannel micropipette. Cells were incubated at 37°C with 5% CO_2_ to promote the formation of embryoid bodies (EBs). On day 2, the medium was changed to a 1:1 mixture of low-FGF and M2 media, followed by 100% M2 on day 3. EBs were cultured in M3 medium from day 4 until day 7. On day 7, EBs were collected, mechanically shredded with a 1000 μL pipette tip, and 3 to 4 shredded EBs were transferred to each well of a Matrigel-coated 6-well plate. Matrigel for coating was diluted 1:30 in DMEM/F-12 and plates were coated for at least 45 min at at 37°C with 5% CO_2_. Shredded EBs were cultured in M3 medium supplemented with ROCK inhibitor (10 µM) and the medium was changed every second day. M2 and M3 media were prepared according to the standard “Basic Media Formulation” without MycoZap. NPCs were passaged 3–4 times to remove remaining EB fragments and generate homogeneous colonies. When NPC cultures reached 80% confluency, they were either subcultured or frozen in Bambanker cryopreservation medium according to the manufacturer instructions. Upon differentiation, cells were fixed with 4% PFA at room temperature for 20 min and characterized using immunohistochemistry staining following the established protocol.

#### Cancer cell culture

GBM14NSC cells were maintained as spheres in stem cell medium containing DMEM (cat. No. 10313021) supplemented with 1xB27 (cat. No. 17504-044), 5% (vol/vol) P/S, 10 ng/mL hEGF, and 10 ng/mL hFGF. To passage GBM14NSC cells, spheres were collected using a 5 ml serological pipette and transferred into a 15 ml conical tube, followed by centrifugation at 300 g for 5 minutes. Upon centrifugation, the medium was carefully aspirated and 1 ml of Accutase solution was added to the cell pellet. The mixture was incubated at 37°C with 5% CO_2_ for 4 minutes to facilitate effective cell dissociation. After incubation, 5 ml of DMEM/F12 was added to the cells to deactivate the Accutase. Cell suspension was centrifuged again at 300 xg for 5 minutes at room temperature. The cell pellet was then resuspended in DMEM/F12, and the appropriate volume of the cell suspension was dispensed into the new flasks.

LN-229 cells were cultured as adherent monolayer, using 6 well plates (Corning, cat. No. 353046), in DMEM (cat. No. 10313021) supplemented with 10% FBS and 5% (vol/vol) P/S. To passage LN229 cells, the culture medium was discarded, and the cell monolayer was rinsed once with PBS 1X. Then 1 ml of 0.25% Trypsin-EDTA solution was added to a well of a 6-well plate and cells were observed under an inverted microscope until the cell layer was fully dispersed, which typically occurred within 2 minutes. Upon incubation, 2 ml of complete growth medium was added to neutralize the trypsin, and cells were distributed to new culture wells. All cell cultures were maintained at 37°C in a humidified environment with 5% CO_2_.

#### Generation of GFP labelled LN229 cells

This cell line was generated by lipofection of GFP-encoding PiggyBac transposon with appropriate transposase plasmid. Τo perform lipofection of LN229 cell line, cells were seeded at a density of 2 × 10^5^ cells per well, in a 6-well plate until they reach 70 – 80% confluency. The lipofection mixture was prepared by combining lipofection reagent and plasmid DNA according to manufacturer’s protocol, using a ratio of 1:3 and a total of 25 μg of DNA. For transfection, the transposase pCAG_hyPBase-SuperPiggyBac and the pB-CAG-GFP vector were used, maintaining a ratio of 2:1 accordingly. The transfection medium was changed after 24 hours at 37°C with 5% CO_2_. To minimize prolonged cytotoxicity by the transfection reagents, cells were washed once with PBS 1X and 1,5 ml of fresh complete medium was added per well. GFP positive cells were sorted in sterile conditions at the Core Facility Flow Cytometry at Biomedical Center Munich of LMU 72h after transfection at single cell density in 96-well plate. Positive clones were selected and subcultured as described above.

Regular PCR tests were conducted in all cultures to check for Mycoplasma contamination according to the kit manufacturer instructions.

#### Generation of iPSC derived monocytes (i-Monocytes)

Monocytes were generated following a modified and optimized version of the protocol by Cao et al. (2020) ^90^, utilizing the HMGU1 iPSC line, the HMGU12 iPSC line, and the HMGU1-RFP iPSC line. Specifically, differentiation of iPSCs into hematopoietic progenitor cells (HPCs) was performed using the STEMdiff Hematopoietic Kit, according to the manufacturer’s instructions. On Days 10 to 12, HPCs were harvested using a 5 mL serological pipette, collected into a 15 mL conical tube and centrifuged at 200 × g for 4 minutes. Cell pellet was washed with sterile 1X PBS and the cells were gently resuspended in 500 μL of IF9S. Cells were seeded at a density of 8 × 10^4^ to 1 × 10^5^ cells per well in a clear flat-bottom Ultra-Low Attachment 24-well plate, with 1 mL of IF9S medium supplemented freshly with 50 ng/mL IL-6, 10 ng/mL IL-3, and 80 ng/mL M-CSF for the induction of monocytes. Two days after i-monocyte induction, 1 mL of fresh IF9S complete media was added. After 5 to 7 days, monocytes were gently collected using a 1000 µL pipette for downstream analysis. For immunofluorescence (IF) analysis, cells were plated on freshly coated coverslips with Geltrex and incubated overnight at 37°C with 5% CO_2_. Upon attachment, the cells were fixed with 4% paraformaldehyde (PFA) for 15 minutes at room temperature and washed three times with 1X PBS for 10 minutes.

#### Glioblastoma multiforme patient selection

Glioblastoma patients were selected at the Department of Neurosurgery at Klinikum Rechts der Isar, Technical University of Munich according to the institutional ethic vote. All adult male patients aged 45 to 75, diagnosed with high-grade glioma, underwent their first surgical intervention for primary glioblastoma, with no history of tumor recurrence. These patients provided preoperative informed consent to participate in the study, which was approved by the relevant ethics committee. Human GB tissue from the peritumoral area was obtained during surgery and processed as described in the Method Details section.

### METHOD DETAILS

#### Protocol and timing for generation of hc-assembloids

##### Day 0: Formation of Embryoid bodies (Ebs); Timing 45 minutes – 1 hour

i. Culture iPSC colonies in a six-well plate until they reach 70–80% confluence. In most cases, one confluent well is enough cells for an entire 96-well plate of Ebs. Prior to Ebs formation the pluripotency of iPSC colonies should be assessed. *<u>Note:</u>* The morphology of stem cell colonies is essential for successful cerebral tissue formation. The colonies should exhibit no signs of differentiation and should demonstrate optimal characteristics indicative of pluripotency. Aside from their differentiation capacity, pluripotency of iPSCs can be assessed with Alkaline phosphatase staining.
ii. Wash iPSCs with PBS 1X in 1 – 2 wells of a 6-well plate. Add 1 ml of pre-warmed Accutase and incubate for 6 – 7 minutes at 37 °C with 5% CO_2_.
iii. Neutralize Accutase with 2 ml of low-FGF medium, gently pipette with 2 ml serological pipette to dissociate cells, and collect the suspension in a conical 15 ml tube.
iv. Centrifuge at 150 xg for 4 minutes at room temperature.
v. Discard the supernatant and gently resuspend the pellet in low-FGF medium supplemented with RI Y27632 (1:200; e.g., 2.5 µl RI in 500 µl medium). Evaluate the cell numbers using a hemocytometer or an automated cell counting device.
vi. Plate 9,000 cells per well in 150 µl of complete low-FGF medium in a low-attachment 96-well U-bottom plate using the multichannel micropipette. Incubate at 37 °C with 5% CO_2_.

##### Day 3: Maintenance of Ebs; Timing 30 minutes

i. Assess the morphology of the Ebs using an EVOS microscope. Small, round Ebs with distinct borders should be visible, although a significant number of dead cells may surround the area around the Ebs (**Fig. 1b**).
ii. Employ a multichannel pipette set to 75-85 μl and pipet gently the medium of the Ebs up and down to remove any debris or cells that may have undergone spontaneous differentiation. *Note:* The pipette should be held vertically, and care should be taken to avoid contacting the Ebs or the bottom of the well.
iii. Tilt the plate slightly to facilitate the settling of Ebs at the bottom, then aspirate approximately 75% of the old low-FGF medium. Replace it with 140 μl of freshly prepared low-FGF medium, without FGF or RI. Maintain the Ebs in the tissue culture incubator at 37 °C with 5% CO_2_.

##### Day 5 – Day 13: Neuronal Induction and Regional Patterning of Ebs; Timing 30 minutes

i. Assess the morphology of the Ebs using an EVOS microscope to follow the progression of Ebs growth and development (**Fig. 1b-b′′′**).
ii. Prepare NIM medium freshly before use. To achieve dorsal and ventral identity patterning of the Ebs, the protocol of Ormel et al. ^28^ was adapted. A drug-patterning treatment was implemented during the neural induction phase from Day 5 to Day 13, with medium changes every two days (days 5, 7, 9, 11). For dorsal identity, use NIM medium without any additions; for ventral identity, add SAG at a final concentration of 0.1 µM along with IWP-2 at a final concentration of 2.5 µM.
iii. To change medium (days 5, 7, 9, 11), replace 75% of the existing medium with the NIM medium containing the appropriate patterning molecules as specified above. Continue the culture of EB at 37 °C with 5% CO_2_ *<u>Note:</u>* Evaluate the morphology of Ebs and neuronal induction before medium changes. Healthy cell aggregates should display smooth edges, with neuroepithelium developing on the outer surface and exhibiting a degree of optical translucency and brightness around the periphery, indicating successful neuroectodermal differentiation. As differentiation progresses, these areas should begin to demonstrate radial organization, resembling a pseudostratified epithelium (**Fig. 1b’’, b’’’**). *Note:* To further promote the growth and development of the newly expanded neuroepithelia, EBs should be transferred to Matrigel without delay, as late transfers may adversely affect the subsequent morphology of cerebral tissues.

##### Day 13: Fusion of Dorsal and Ventral EBs and Embedding in Matrigel; Timing 1 – 2 hours

i. Thaw Matrigel overnight on ice (place the ice container at 4^0^C) prior to use.
ii. Use sterile scissors to cut a 200-µl pipette tip, creating an opening of 1–1.5 mm in diameter.
iii. Create the parafilm wells by pressing a piece of parafilm onto the surface of a sterile Terasaki plate. This will serve as the base for embedding the EBs. Using disposable forceps at a 45° angle, press down into the Parafilm at each hole in the tip tray to form small wells for holding the EBs.
iv. Carefully transfer EBs from each well of the 96-well plate to culture dishes containing 5 ml of IDM-A. Ensure that each dish corresponds to the patterned EBs, resulting in two distinct dishes: one for dorsal and one for ventral EBs.
v. Transfer one dorsal EB and one ventral EB into the same Parafilm well. Carefully remove as much medium as possible without aspirating the EBs. Add a single drop (∼30 µL) of Matrigel to the well using a cold P200 pipette tip, taking care to avoid the formation of bubbles. Ensure that there are no delays to maintain the integrity of the Matrigel.
vi. Examine the position of the EBs using the stereoscope and gently position the two Ebs close together in the center of the Matrigel droplet, using a 200 µL pipette tip.
vii. Transfer the parafilm sheet containing the EBs to a 100 mm cell culture dish. Incubate at 37 °C for at least 45 minutes to allow the Matrigel to fully solidify.
viii. After solidification, gently add 25 ml of IDM-A to each 100 mm culture dish, using a 25 ml serological pipette, to wash the EBs off the parafilm, taking care not to disturb the newly embedded assembloids.
ix. Place the embedded EBs in the incubator maintained at 37 °C with 5% CO_2_ for 3 days.

##### Day 17: Assessment of hc-assembloid formation and long term culture; Timing depends on the number of Ebs

i. Monitor efficient fusion using the EVOS microscope. Tissues should begin forming buds of more expanded neuroepithelium containing fluid-filled cavities (**Fig. 1b′′′**). The fusion process is typically efficient and should occur within one week if the EBs are positioned sufficiently close to each other. If any visible spaces remain between hc-assembloids after one week, these “failed” fusions can be removed by aspiration. *<u>Note:</u>* Frequently, more migratory cell types disperse from the main assemboid into the remaining Matrigel droplet. These cells are fibroblast-like and likely represent a small subset of non-neural identities that evaded neural induction and they do not survive in IDM medium while their movement away from the tissue enhances the development of neuroepithelial buds.
ii. Tilt the dish to change the medium, allowing the droplets to settle, and carefully aspirate as much of the medium as possible without disturbing the hc-assembloids.
iii. Replace medium with 20ml fresh IDM+A and move the fused hc-assembloids to an orbital shaker (55 rpm).

##### Day 17-104: Long term culture and growth of hc-assembloids

Replace medium every 5 days with fresh IDM+A. After day 60, change medium with fresh IDM+A every 2-3 days as described above. Continue the culture of hc-assembloids at 37 °C with 5% CO_2_.

##### Day 104-110: Transplantation of cancer cells or NPCs to hc-assembloids using a microinjector; Timing depends on number of hc-assembloids

i. Set-up the transplantation station. Set the biosafety hood to cleaning mode and raise the safety glass to assembly the transplantation station. Spray the stereoscope with 80% ethanol and position it on the left side of the hood. Adjust the stereoscope to ensure a clear, magnified view of the samples, with appropriate working distance and alignment with the microinjector. Clean the microinjector with 80% ethanol, then place it on the right side of the hood. Confirm that the microinjector is properly calibrated, securely mounted, and capable of precise control for transplantation. Ensure both the stereoscope and microinjector are stable, easily accessible, and that there is sufficient space in the hood for smooth operation throughout the procedure. Utilize 100 × 0.9 mm^2^ glass capillary needles fabricated using a puller, with a ‘one-stage’ pull setting at 63.5 °C and sterilized under UV before use. Spray with ethanol and place inside the hood a pair of metallic forceps, culture dishes P/S 100/20 mm, tapered tips for loading the glass capillaries and a piece of parafilm.
ii. To prepare cancer cells prior to transplantation, culture LN229, GBM14NSC cells and NPCs until they reach 80% confluency.
iii. Prepare a single-cell suspension for each cell line in organoid culture media (IDM+A) at a concentration of 2,000 cells/µl as described previously for cancer cells and a single cell suspension at a concentration of 20,000 cells/μl for NPCs.
iv. Keep the cell suspensions on ice during the transplantation procedure to preserve cell viability.
v. Examine the hc-assembloids under the stereomicroscope to confirm their integrity.
vi. Use a sterile surgical scalpel to cut the end of a 1000 µL sterile pipette tip. Then, using a 1000 µL micropipette, carefully transfer the hc-assembloids into a sterile 100 mm culture dish.
vii. Position each hc-assembloid inside a droplet of IDM+A medium to immobilize them for the transplantation procedure.
viii. Gently aspirate 10 µL of the prepared cancer cell suspension using a tapered pipette tip.
ix. Load the cell suspension into a sterilized, pulled and sterilized 100 × 0.9 mm^2^ glass capillary needle.
x. Using metallic forceps, cut the sealed end of the glass capillary under the stereomicroscope to allow cell release during injection.
xi. Before proceeding with transplantation, release a droplet of the cancer cell suspension onto a piece of parafilm to ensure that the injection volume is exactly 1 µl, corresponding to 2,000 cells for cancer cells and 20,000 cells for NPCs.
xii. If the injection volume is not accurate, adjust the settings of the microinjector by modifying the injection time and precure pressure to achieve the correct volume.
xiii. Gently insert the capillary needle into the desired site of the organoid, preferably near the periphery to avoid injection of cells in the necrotic areas. Avoid penetrating the organoid too deeply to prevent structural damage.
xiv. Release 1 µL of the cancer cell suspension, ensuring not to disrupt the organoid structure. After the injection is complete, withdraw the needle gently.
xv. Repeat the procedure for additional hc-assembloids as needed.
xvi. Add 25 mL of fresh IDM+A medium to each plate, taking care not to disturb the transplanted hc-assembloids.
xvii. Place the plates in the incubator at 37°C with 5% CO_2_ for 24 hours without shaking them.
xviii. Transfer transplanted hc-assembloids onto an orbital shaker set at 55 RPM for optimal growth and nutrient distribution 24h after transplantation.
xix. Change the medium every 3-5 days, carefully aspirating the old medium without disturbing the hc-assembloids.

##### Troublshooting

###### Problem 1

Low Cell Viability

###### Potential solution

Ensure that Accutase incubation time is optimized for gentle detachment. Avoid excessive incubation to prevent cell damage. After detachment, resuspend cells carefully in low FGF media supplemented with RI to enhance viability. If necessary, adjust Accutase incubation time based on cell behavior.

###### Problem 2

EB Disruption

###### Potential solution

Use low-attachment plates to support EB formation and minimize disruption. Pipette gently to prevent shear stress, which can disaggregate Ebs. Avoid excessive mechanical handling.

###### Problem 3

Failed Neuronal Induction

###### Potential solution

Assess iPSC health before differentiation to confirm the starting population is viable. Verify accurate cell counting and consistently plate 9,000 cells per well. Ensure uniform distribution of small molecules in the media to promote differentiation.

###### Problem 4

Inconsistent Patterning

###### Potential solution

Check the activity and concentration of key small molecules such as SAG and IWP-2. Prepare fresh stocks if necessary. Replace media at precise intervals to maintain consistent signaling gradients for patterning.

###### Problem 5

Failure of Matrigel to Solidify during Embedding of Ebs

###### Potential solution

Thaw Matrigel properly and keep it on ice until use. Ensure the incubation time is sufficient— at least 45 minutes—for solidification. Avoid temperature fluctuations during preparation.

###### Problem 6

Inadequate hc-Assembloid Fusion

###### Potential solution

Carefully position dorsal and ventral Ebs close together within the Matrigel droplet to enhance fusion. Minimize movement post-placement to prevent separation.

###### Problem 7

Inaccurate Injection Volume

###### Potential solution

Before transplantation, release a droplet of the cell suspension onto parafilm to confirm the injection volume is precisely 1 µl. Adjust the injection time and precure pressure settings on the microinjector if necessary.

###### Problem 8

Clogging of the Glass Capillary Needle

###### Potential solution

Load the capillary slowly to prevent air bubble formation. Remove the loading tip carefully to avoid introducing bubbles. If clogging persists, check for cell aggregation and gently resuspend before reloading.

###### Problem 9

No Evidence of Cancer Cells Post-Transplantation

###### Potential solution

Maintain the cancer cell suspension on ice throughout the procedure to preserve viability. Avoid prolonged exposure to room temperature or mechanical stress during transplantation.

###### Problem 10

Damage to hc-Assembloid Structure During Injection

###### Potential solution

Ensure gentle insertion of the needle, avoiding deep penetration into the organoid. Inject slowly to minimize tissue disruption. Target peripheral areas for injection, maintaining pressure between 70-85 psi.

###### Fixation and cryosectioning of hc-assembloids

i. Transfer hc-assembloids from culture dish to a 15 mL conical tube using a 1000 ul micropipette with a cut sterile tip to avoid damaging the hc-assembloids.
ii. Rinse the hc-assembloids gently with PBS 1X to remove any remaining culture medium.
iii. Add 8 mL of freshly prepared 4% PFA to the tube containing the hc-assembloids
iv. Incubate at 4°C overnight. Ensure that the hc-assembloids are fully submerged in PFA to allow even fixation.
v. Wash the hc-assembloids with 10 ml PBS 1X (3 times, 15 minutes) to remove residual PFA.
vi. Add 10 ml of 30% sucrose solution after washing in PBS 1X.
vii. Incubate the hc-assembloids in sucrose at 4°C without agitation until they sink to the bottom of the tube, indicating complete saturation. This may take 24 – 48 hours depending on the size of the tissue
viii. Ensure that the hc-assembloids are fully submerged in the sucrose solution to prevent damage during freezing.
ix. Prepare cryomolds by filling them halfway with Neg-50^TM^ embedding compound.
x. Carefully transfer the sucrose-treated hc-assembloids from the sucrose solution into the cryomolds using a metallic spatula.
xi. Ensure the hc-assembloids are centered in the Neg-50^TM^ -containing cryomolds.
xii. Fill the cryomold completely with Neg-50^TM^ compound, ensuring no air bubbles are trapped around the hc-assembloids and rapidly place them on dry ice.
xiii. Transfer the frozen cryomolds to a −80°C freezer until cryosectioning
xiv. For sectioning, place the cryomolds in a cryostat pre-cooled to −20°C. Allow the tissue to acclimate to the temperature for 10–15 minutes before sectioning.
xv. Cut sections of the desired thickness (typically 16 µm) and place them on superfrost slides for further staining or analysis.

#### hc-Assembloids Immunohistochemistry

Tissue processing was carried out as described above. For immunolabeling of hc-assembloids, we used the primary antibodies listed in Key Resource Table. Primary antibodies were visualized using specific secondary antibodies conjugated with Alexa Fluor 488, Alexa Fluor 546, or Alexa Fluor 633 (1:1,000, Invitrogen, Thermo Fisher). Tissue sections were embedded in Aqua PolyMount.

#### Image acquisition and processing

All confocal images were obtained using an Olympus FV1000 cLSM system (Olympus) with FW10-ASW 4.0 software, and a Leica TCS SP8 system (Leica) operated with Leica Application Suite X software (v.3.5.6.21594) at the Core Facility Bioimaging, at the Biomedical Center of Ludwig-Maximilians-Universität. All presented images are maximum intensity projection of confocal Z-stack.

#### Whole hc-assembloids clearing with and Immunostaining

Dent’s Fix solution was freshly prepared by combining dimethyl sulfoxide (DMSO) and methanol (MeOH) in a 1:4 ratio. For Dent’s Bleach solution, 30% (vol/vol) hydrogen peroxide (H_2_O_2_) was mixed with Dent’s Fix in a 1:2 ratio. For BABB solution preparation, benzyl alcohol and benzyl benzoate were mixed in a 1:2 ratio. hc-assembloids were fixed in 4% PFA (wt/vol) at 4°C for 3 hours and rinsed three times with PBS 1X. Samples were bleached in Dent’s Bleach solution for 24 hours at 4°C with gentle rotation, followed by 5 rinses with MeOH. After rinsing, samples were fixed in Dent’s Fix solution for 24 hours at 4°C with gentle rotation, rinsed 3 times 10 min and 3 times 1h with 1X PBS at room temperature prior to addition of primary antibody. Primary antibodies in blocking solution (75% PBS 1X, 20% DMSO, and 5% NDS) were incubated for 48 hours at 4°C with rotation. Following 5 washes of 1h with PBS 1X, secondary antibodies conjugated to Alexa Fluor 488, Alexa Fluor 546, or Alexa Fluor 633 were applied for 48 hours at 4°C with rotation. After secondary antibody incubation, samples were washed 5 times with 1X PBS at room temperature for 1 hour each time, followed by overnight wash in 1X PBS at 4°C with rotation. Half of the PBS 1X was replaced with MeOH and incubated for 10 minutes at room temperature. Samples were then washed 3 times with MeOH for 20 minutes each at room temperature. Subsequently, half of the MeOH was replaced with BABB solution and incubated for 10 minutes. Samples were cleared in 100% (vol/vol) BABB solution and embedded in 2% low melting agarose for imaging.

*Note:* The blocking solution was stored at 4°C for maximum 5 days and BABB/methanol and BABB were always freshly prepared.

#### 3D Imaging of cleared hc-assembloids

Cleared hc-assembloids were imaged using Lightsheet Fluorescence Microscopy (LaVision UltraMicroscope II, Blaze, Miltenyi BioTec) and processed with LaVision BioTec ImSpector Software at the Core Facility for Pathology and Tissue Analytics at Helmholtz Zentrum Munich.

#### In situ hybridization

RNA in situ hybridization was performed using RNAscope® Multiplex Fluorescent Reagent Kit according to the manufacturer’s instructions. Briefly, hc-assembloid sections were fixed in 4% PFA at 4 °C for 15 min, ethanol-dehydrated, treated with H_2_O_2_ and protease-permeabilized for 20 min at 40 °C. hc-assembloid sections were incubated for 2 h at 40 °C. Samples from plated GBM14NSC and LN229 cancer cells were treated according to manufacturer’s instructions (User manual Cat. Nr: MK 50-012, Rev A/ Date 092520, ACD). All samples were incubated using the following probes: *SEMA5A*, *NTRK2*. Subsequently, immunohistochemistry analysis for GFP was performed, as described above.

#### scRNA-seq library preparation

Cells were isolated from hc-assembloids coming from following groups: intact, media transplanted, GBM14NSC transplanted and LN229 transplanted (n (hc-assembloids)= 3 per group) from three different biological replicates (independent batches) and dissociated into a single-cell suspension using the Papain Dissociation System, according to manufacturer’s instructions. Incubation in dissociating enzyme was performed for 45 min with titration in the incubator at 37 °C with 5% CO2. GBM14NSC and LN229 cancer cells for scRNAseq were collected with certification at 300 xg for 5 minutes.

Single-cell suspensions were loaded into 10x Genomics Single Cell 3′ Chips. Libraries were prepared according to the manufacturer’s instructions for Chromium Single Cell 3′ Reagent Kits v3.1 (10x Genomics), pooled and sequenced on a NovaSeq6000 (Illumina) with the recommended number of cycles according to Chromium Single Cell 3′ Reagent Kits v3.1, Dual Index: Read 1:28 cycles, i7 index: 10 cycles, i5 index: 10 cycles and Read 2: 90 cycles. Sequencing was performed by the Core Sequencing Facility at the Helmholtz Zentrum Munich and NGS Core Facility, Institute of Human Genetics; Division of Genomics; Life & Brain Research Centre; University Hospital of Bonn.

#### Single-cell RNAseq Analysis

All samples were aligned on the human reference genome GRCh38-2024-A using Cellranger v10.0.0 ^121^ Ambient RNA removal was performed using cellbender 0.3.2 ^122^, the resulting counts were analysed in python 3.11 using Scanpy 1.11.5 ^123^. [https://link.springer.com/article/10.1186/s13059-017-1382-0].

Cells first went through quality controls, preliminary filtering, doublet detection and filtering using scrublet v0.2.3 ^124^. The resulting count matrix was then filtered as followed: minimum count of 1500 UMIs per cell, maximum count of 10000 UMIs per cell, maximum percentage of mitochondrial counts less than 20%, 1000 minimum gene detected per cells, 9000 maximum genes detected per cells. The resulting count matrix of all samples after final filtering contains 134,030 cells x 33,598 transcripts. Batch correction was done with scvi-tools v1.4.1 ^125^,the data was normalized using Scanpy normalize_total function.

After the data processing, and batch correction, the remaining batch effects were measured using scib v1.1.7 ^126^ using the following three metrics:

- Cosine similarity: measures for each cell the gene-expression difference to the reference samples. Values closer to 1 indicate a higher similarity. Values are averaged per cell-types or cell clusters.
- Pearson correlation: evaluates the linear correlations between gene expression across batches per cell-types (or cell clusters).
- Nearest-Neighbor Mixing: for each cell in the neighbourhood graph, it calculates the proportion of a cell’s nearest neighbour originate from a different batch. Higher values represent a bigger batch mixing. The values were averaged per cell-types or cell clusters.

The batch effect evaluation was performed per cell type on hc-assembloid derived cells to assess the batch effect of media transplanted sample compared to an intact hc-assembloid and the batch effect of GB transplantation. A second batch effect was assessed, to test whether the change of media of GB cells from a growth media to an organoid media. To this end, all the GBM14NSC samples (cells and transplanted organoids) were processed together. Organoid derived cells were excluded before the batch effect evaluation.

Differential gene expression was performed using non-parametric Wilcoxon Rank Sum test with Benjamin-Hochberg multiple hypothesis testing correction. Differential genes were defined as having an adjusted p-value less than 0.05 with a log2 fold change greater than 1. Subsequent genes were used for functional enrichment analysis using gprofiler-official v1.0.0 ^127^. To this end, differential genes were compared to a custom background represented by all genes used for the Wilcoxon Rank Sum test. The enrichment was restricted to Gene Ontology Biological Process terms, and the GO term enrichment was corrected using Bonferroni multiple testing correction method. Finally, significant GO terms were filtered out to contain at least 1% of the genes in the query list and an adjusted pvalues < 0.05.

Following the same scRNA-seq processing approach as mentioned above, all intact hc-assembloid samples were processed together to annotate the different brain hc-assembloid cell types present in the data. Preliminary cell labels were established using known cell markers as described in Uzuiano et al. ^33^ and Velmeshev et al. ^128^. Cells were clustered using leiden and cell-type scores were calculated using Scanpy ^123^ leiden and score_genes functions. The cell labels were confirmed and refined using differential gene expression and functional enrichment analysis to obtain final detailed cell labels. Cell-cycle scores were the score_gene function, using the KEGG cell cycle genes from bHarmonizome platform ^43,44^. All marker genes used for cell cycle scoring are available in Table S10.

The cell annotation was transferred from the intact hc-assembloid object to the processed object containing all samples. Then, the cell label was extended to all hc-assembloid derived cells based on a majority voting approach. After a detailed leiden cluster, cell label was assigned as the majority cell label coming from the intact hc-assembloid cells.

To determine shifts in glia cluster composition upon GB transplantation, the data was first subset to only consider glial cells coming from GB transplanted or media transplanted samples. Then, compositional analysis was performed using a generalized linear model assuming a binomial distribution and using the media transplanted cells as reference. To accounts for zero cell count in some clusters, a Haldane-Anscombe correction was applied, and a Bonferroni was applied for multiple testing correction. Finally, differential clusters were identifies as having an adjusted p-value < 0.05 and ranked based on log odds ratios.

To identify a gene signature specific to radial glia cells after GB transplantation, differential gene expression analysis was performed for all significant enriched clusters (log odds ratios >0) against depleted clusters (log odds ratios <0). All differential genes with an adjusted p-value <0.05 and a log2 fold change >0 were considered. The intersect of genes identified as differential for all enriched clusters are used to define the glial reactive signature.

#### Cell-cell-communication Analysis

To identify cell-type specific cell-cell communication specific to cells after GB transplantation, cell-cell communication has been both measured in a control sample set, containing media transplanted samples and GB cells, and GB transplanted sets, either LN229 or GB14NSC transplanted samples. We rely on the cellphoneDB ^70^ cell-cell communication algorithm as implemented in LIANA+ ^129^ v0.1.9. The primary cell-cell communication analysis is restricted to cell clusters identified as derived from the GB cells or labelled glial cells are inputted in cellphoneDB algorithm.

To identify meaningful cell-cell communication dynamics arising upon GB transplantation, while accounting for cell composition biases, a custom cell-type specificity normalization was implemented. First, the p-values are transformed into specificity scores defined as (1 - pvalue) to emphasize high-confidence Ligand-Receptor interactions. Then the differential specificity is calculated by subtracting GB transplanted and control set interaction scores to highlight novel or differential interactions upon GB transplantations, independently of cell cluster abundance. Furthermore, only interactions with a pvalue <0.05 within the GB cancer set are considered further. Finally, to reduce the importance of ubiquitous Ligand-Receptor pairs across the dataset, which otherwise may dominate the cell-cell communication analysis, each Ligand-Pair receptor interaction per cell type was divided by the average expression of the pair across all cell clusters across the datasets.

A second cell-cell communication analysis was performed, to quantify the interaction strength between cancer cells and hc-assembloid derived cells with either dorsal or ventral identity. To this end, the cellphoneDB algorithm was run on a subset of the GB transplanted samples, which was restricted to GB cells and hc-assembloid derived cells with dorsal and ventral identities. The overall cell-interaction strength between clusters was calculated as the average expression levels of the Ligand-Receptor pairs.

Dorsal and ventral scores are calculated using Scanpy ^123^ score_genes function using gene markers as specified in **Table S1**. Cells with positive scores were labeled as either dorsal or ventral. In the case were both dorsal and ventral scores were positive, only the maximum score was considered.

#### Spatial data Analysis

The processed data was obtained on Zenodo from Greenwald et al. ^80^. The data was loaded and analyzed using Squidpy v1.8.3 ^130^. Malignancy scores are calculated based on malignant metaprograms determined by Greenwald et al. ^80^. Both malignancy and reactivity scores are determined using scanpy sc.tl.score_genes function. Spatial organization between malignancy and reactivity scores was assessed using Squidpy Moran’s I index to quantify the spatial patterning and Spearman correlation to assess cross-signature correlations.

#### Flow cytometric analysis of i-monocytes

For flow cytometric analysis, cells were incubated with fluorescent-conjugated antibodies against CD14 and CD16 diluted 1: 50 in PBS 1X at 4°C for 30 minutes in the dark, washed twice with 1X PBS and analyzed at the Core Facility Flow Cytometry at the Biomedical Center Munich of LMU using a BD Fortessa (5-Laser) cytometer. The FACS gates were set using cells incubated with fluorescently labelled isoform control antibody (see Key Resource Table) diluted in PBS 1X.

#### Inoculation of i-Monocytes with hc-assembloids

For inoculation of i-monocytes to hc-assembloids, one hc-assembloids was added per well of the Ultra-Low Attachment 24-well plate containing i-monocytes and incubated for 48 hours at 37°C with 5% CO_2_ without shaking. After the incubation period, the hc-assembloids were transferred back to a 100 mm culture dish containing IDM+A and incubated at 37°C with, 5% CO_2_ with shaking for a total of 5 to 7 days, after which the hc-assembloids were harvested for downstream experimental procedures.

#### Isolation of human peripheral blood mononuclear cells (PBMC)

The PBMCs were isolated from healthy human donors blood using Ficoll gradient according to manufacturer’s instructions.

#### Collection and Preparation of Human Tissue Samples

Tissue samples were collected from glioblastoma patients during surgery and immediately fixed in 4% PFA overnight at 4°C for tissue preservation. Following fixation, tissues were transferred to 70% isopropanol for a brief wash before undergoing dehydration through an ascending alcohol series: 70%, 80%, 90%, and 100% isopropanol (each for 10-30 minutes). Samples were incubated overnight with 100% isopropanol to ensure sufficient dehydration. The tissues were then cleared with two changes of xylene for 35 minutes each, ensuring complete dehydration and solvent removal. After xylene treatment, the samples were embedded in paraffin using grass glass containers at 56°C and incubated in paraffin at 55°C for 2-5 days to ensure full embedding. The paraffin blocks were then cast on metal rails, allowed to cool to room temperature, and detached for subsequent processing. Paraffin blocks were cut using a microtome in 6-8 μm thin sections as described previously ^131^.

#### Immunofluorescence Staining of Formalin-fixed paraffin-embedded (FFPE) Human Tissue Sections

Formalin-fixed paraffin-embedded (FFPE) human tissue sections were deparaffinized in xylene (2 × 3 minutes), rehydrated through 100% isopropanol (2 × 3 minutes) and a graded isopropanol series (90%, 80%, 70%; 3 minutes each), and washed in PBS 1X for 10 min. Sections were permeabilized and blocked in 10% NGS in 0.3% Triton X-100-PBS 1X for 30 min at room temperature. Lipofuscin autofluorescence was quenched using TrueBlack® for 1 min, followed by PBS 1X washes. Primary antibodies were incubated overnight at 4°C in PBS1X containing 10% NGS. The following day, sections were washed in PBS 1X three times 15 minuteseach and incubated with fluorophore-conjugated secondary antibodies for 2 h at room temperature light protected. Nuclei were counterstained with DAPI for 10 min, followed by three PBS1X washes of 15 minutes each. Sections were mounted with Aqua Poly/mount and dried overnight at room temperature in the dark prior to imaging.

#### Generation and Maintenance of Primary Ninkovic Glioblastoma Lines (NGL)

Primary glioblastoma (GB) cell lines were derived from high-grade glioma specimens obtained during surgical resection under sterile conditions. Fresh tumor explants were transferred into stem cell medium containing DMEM (cat. no. 10313021) supplemented with 1xB27 (cat. no. 17504-044), 5% (vol/vol) P/S, 10 ng/mL hEGF, and 10 ng/mL hFGF and maintained on ice until processing. Tumors were mechanically minced into small pieces and enzymatically digested using the Papain Dissociation System (Worthington Biochemical Corporation, LK003150) according to the manufacturer’s instructions. Briefly, tissue fragments were incubated in papain solution for 45-90 min at 37°C with gentle trituration followed by filtration and centrifugation. Cells were resuspended in complete stem cell DMEM/F12 medium and seeded onto Geltrex-coated T25 flasks.

The following day, non-adherent tumor cells were harvested and transferred to uncoated T25 flasks for expansion as floating tumorspheres in GB stem cell medium supplemented with EGF and FGF. Cultures were maintained at 37°C and 5% CO_2_, with medium change every 4 to 5 days.

#### PKH26 Labeling of NGL Cells

NGL cells were labeled using the PKH26 Red Fluorescent Cell Linker Kit (Merck, PKH26GL-1KT) according to the manufacturer’s instructions. Cells were harvested at 80% confluency, washed with PBS1X to remove serum proteins, and resuspended in Diluent C, at ∼1 × 10⁶ cells/ml. PKH26 dye was diluted in Diluent C immediately before use and mixed with the cell suspension for 5 min at room temperature light protected. Labeling reaction was quenched by addition of complete medium containing 10% FBS, followed by three washing steps with centrifugation at 300 × g for 5 min to remove excess dye. Labeled cells were resuspended in GB stem cell medium and maintained on ice until downstream processing. Labeling efficiency and cell viability were assessed by fluorescence microscopy prior to experiments.

#### Isolation of Extracellular Vesicles (EVs) from culture medium

Extracellular vesicles (EVs) were isolated from conditioned culture medium collected from hc-assembloid cultures, using an adapted protocol from D’Acunzo et al. (2017). Briefly, 25 ml of medium was collected and subsequently centrifuged at 300 × g and 2,000 × g at 4°C to remove dying cells and debris, followed by filtration through a 0.22 μm filter. Cleared samples were snap frozen in liquid nitrogen and stored at −80°C until further processing.

For EV isolation, samples were thawed at 4°C overnight and subjected to ultracentrifugation at 10,000 × g for 30 min. The pellet was discarded and supernatant was processed by ultracentrifugation at 100,000 × g for 70 min at 4°C. EV enriched pellets were resuspended in sterile cold PBS 1X without Ca^2^⁺/Mg^2^⁺ and used for downstream nanoparticle tracking analysis (NTA). For NTA measurements, EV preparations were diluted further 1:1000 in PBS 1X prior to analysis.

#### Electron Microscopy imaging of EVs

EV samples were incubated on formvar/carbon-coated glow-discharged copper grids for 20 min. Grids were washed two times with PBS and subsequently fixed by incubating the grid with glutaraldehyde in PBS for 5 min. Grids were washed with water and stained with neutral uranyl acetate for 5 min and coated with methylcellulose/uranyl acetate for 5 min. Grids were air-dried at room temperature and analyzed using a JEM-120i transmission electron microscope (JEOL Nordic AB) operated at 120 keV. Images were captured by a Qedira CMOS camera (Emsis, GmbH, Germany).

#### Nanoparticle Tracking analysis

Prior to measurement, the NTA instrument was calibrated according to the manufacturer’s instructions, and the sample chamber was rinsed with distilled water. A calibration nanobead solution was prepared by diluting polystyrene beads 1:200,000 in Milli-Q water. Subsequently, 20 ml of the diluted calibration solution was loaded to the instrument to verify accurate system performance and optimize particle detection settings. EV samples were diluted in sterile PBS 1X to achieve a particle concentration within the optimal detection range of the instrument. For each measurement, 1 ml of the diluted EV suspension was loaded into a 1.5 ml syringe and injected into the sample chamber. Measurements were performed under constant flow conditions and recorded using identical camera and detection settings for all samples to ensure comparability. Between measurements, the chamber was thoroughly rinsed with Milli-Q water to prevent sample cross-contamination. Data were analyzed by nanoparticle tracking analysis using a ZetaView instrument (S/N 22-847; Particle Metrix, Germany) and ZetaView software (version 8.06.01 SP1).

#### Blocking EV biogenesis and trafficking

For inhibition of EV biogenesis, transplanted hc-assembloids were cultured in IDM+A with 5 µM GW4869 (Merck, D1692) for the final week of culture, from 2.5 to 3.5 weeks post-transplantation. The conditioned culture medium, supplemented with GW4869 was replaced daily throughout the treatment period. At the end of treatment hc-assembloids were fixed for downstream analysis. For blocking EV trafficking, transplanted hc-assembloids were cultured in IDM+A supplemented with 10 µM Y-27632 (Millipore, SCM075) for 1 week, from 2.5 to 3.5 weeks post-transplantation. Conditioned culture medium was refreshed daily throughout the treatment period. Conditioned media were collected 24 hours after treatment initiation and again at the end of the treatment period for EV isolation as described above. Following 7 days of Y-27632 treatment, hc-assembloids were fixed for downstream analysis.

#### EV Fluorescent Labelling with PKH26

Isolated EVs were fluorescently labeled using the PKH26 membrane using a modified version of the manufacturer’s protocol. The dye working solution was prepared by diluting 6 µl of PKH26 dye in 1 ml of Diluent C. The EV suspension was prepared by mixing 20 ul of the prepared EV stocks solution (1 × 10^7^ EVs/μl) with 80 ul of Diluent C. To initiate labelling 100 ul of the PKH26 working solution were added to the EV solution. The mixture was incubated at room temperature for 5 minutes, with gentle pipetting every 1 minute to ensure uniform EV labelling. To terminate the labelling reaction, 2 ml of 10% bovine serum albumin (BSA) (Merck, cat. no. A1595) in PBS was added to the mixture and the sample volume was then adjusted to 10 ml with PBS1X without calcium (Ca^+2^) and magnesium (Mg^+2^). Unincorporated PKH26 dye was removed from labeled EVs by ultracentrifugation and the EV-containing pellet was resuspended in PBS1X at a volume of 20 μl to preserve initial concentration of 1 × 10^7^ EVs/μl. Upon labelling intact hc-assembloids were injected with 1 ul of PKH26 labelled EVs from media collected from cultures of hc-assembloids, LN229 cultured cells and LN229 transplanted hc-assembloids. Transplantation was performed as described in the corresponding section with 1 ul per hc-assembloid.

#### Quantification of mechanical stiffness in hc-assembloids

The effective Young’s modulus (stiffness) was quantified inside tumor and tumor free regions using magnetic droplets as force actuators ^59^ as recently described ^60^. Controlled stresses were applied within the tissue for durations ranging from 10 to 45 minutes using magnetic droplets injected into hc-assembloids four hours prior. The strain response of the tissue was quantified through shape analysis of the fluorescently labeled magnetic droplet, which was captured with time series confocal imaging (timestep: 1.5 to 4 seconds). The strain was extracted via fitting ellipsoids to the deformed droplet-tissue boundary. Droplet pre-deformations were considered using Mohr’s geometric representation of strain ellipses ^60^. To quantify the short-term elastic response, we fit the strain curve with a five-parameter generalized Maxwell solid (GMS) model ^59^. The model consists of 3 parallel branches: one spring (*k*_0_) and two maxwell elements represented by a spring (*k*_1_ and *k*_2_, respectively) in series with a dashpot. The magnitude and orientation of applied stress was determined by calibration in N190000 Viscosity Standard (Cannon Instrument Company) at 37 °C. We recently demonstrated that long-term tissue stiffness dominates over droplet capillary stresses in other neuronal organoid tissues ^60^. Thus, we report the short-term Young’s modulus as *E* = (5/12)(*k*_0_ + *k*_1_ + *k*_2_), where *k*0 + *k*_1_ + *k*_2_ is the sum of the spring constants in the GMS model and *k*_0_ represents long-term residual tissue stiffness, not droplet capillary stiffnesses.

#### Sliced hc-assembloid culture

##### Live imaging of grafted cerebral hc-assembloids

Following two weeks post transplantation hc-assembloids were embedded in 3% low melting agarose in DMEM/F12 and sectioned into 300 μm slices with a Leica VT1200S vibratome. The hc-assembloid slices were cultured free floating in IDM+A media in 12-well plates, on an orbital shaker at 37°C 5% CO2, for 2 weeks. One day prior to imaging, the hc-assembloid slices were stained with BioTracker 655 Red Cytoplasmic Membrane Dye (1 μl/mL, in IDM+A) overnight on the orbital shaker. Upon incubation hc-assembloids with were embedded and immobilized in a Matrigel droplet on an ibidi sterile and uncoated 35mm high #1.5 glass bottom µ-Dish (Cat. 81218-200) with 2mL of culture medium. The preparation was placed onto an ASImaging plate holder (Cat.I-3091) and incubated on a microscope-mounted EMBL Incubator GP169 maintaining 37°C and 5% CO2. 24h live imaging was performed using a Z1 Observer spinning disk microscope (Zeiss) equipped with a Yokogawa CSU-X1 scanner unit and an Evolve Delta Photometrics EMCCD camera (512 x 512 pixels FOV). The acquisition was performed with a 20x LD Plan-NEOFLUAR NA 0.4 PH2 Korr objective (Zeiss Cat. 421351-9970). 150 μm Z-stacks were captured at a 5μm interval and maximum projections were produced. Each position and channel were acquired every 15 minutes. GFP expressing cells were exposed to 5% laser at 100ms exposure time and RFP expressing cells were exposed to 3% laser at 100ms exposure time. Both channels were acquired consecutively. The pre-processing was done on the Zeiss ZEN 3.2 software (3.2.0.010 build). Videos were assembled and optimized using Fiji ImageJ, with maximum projections, median filtering, stack registration and rotations applied.

#### Generation of microglia-free cerebral organoids

Cerebral organoids were generated following Lancaster et al. 2014 directions with some small changes ^27^. 24 hours prior EB generation, iPSCs were treated with 1% DMSO in mTeSR1 media. iPSCs were dissociated into single cells StemPro Accutase Cell Dissociation Reagent (Life Technologies, A1110501) and plated in the concentration of 2000 single iPSCs/well into low-attachment 96-well tissue culture plates in mTeSR medium supplementer with Rock inhibitor (50 μM final concentration). After 48 hours, media was replaced with Forebrain medium 1 (DMEM/F12 supplemented with 20% Knockout Serum Replacement, 1% nonessential amino acids, 1x GlutaMAX supplement, 0.1 mM 2-mercaptoethanol and 1x antibiotic-antimycotic) supplemented with 100 ng/ml Noggin (StemCell, 78060). Media was replaced any second day and on day 5, media was replaced with NIM medium supplemented with 100 ng/ml Noggin. On day 7, EBs were embedded in Matrigel Cookies as explained in Qian et al., 2018 ^132^. On day 9, media was replaced to IDM-A (DMEM/F12-GlutaMAX and Neurobasal in ratio 1:1 supplemented with 1:200 N2 supplement 1:50 B27 without vitamin A, 0.5% nonessential amino acids, insulin 2.5 μg/ml, 1:100 Antibiotic-Antimycotic, and 50 μM 2-mercaptoethanol) and changed every second day. On day 14, Matrigel cookie was removed and media was replaced by IDM+A (DMEM/F12-GlutaMAX and Neurobasal in ratio 1:1 supplemented with 1:200 N2 supplement 1:50 B27 with vitamin A, 0.5% nonessential amino acids, insulin 1 μg/mL, 1:100 antibiotic-antimycotic, and 50 μM 2-mercaptoethanol, 1 mg/ml NaHCO3) and organoids were transferred onto the orbital shaker and medium was replaced twice per week.

#### Quantification and statistical analysis

Quantification of area coverage fluorescence intensity was performed using confocal immunofluorescence z-stack images analyzed by ImageJ. Specific regions of interest (ROIs) were defined of 250 x 250 μm to ensure consistent measurements across all samples. Area coverage of marker positive signal was determined by thresholding fluorescence intensity in ‘Max Intensity’ projections, following background subtraction. Quantification of signal fluorescence intensity was performed using confocal immunofluorescence z-stack images analyzed in ImageJ. For each image, specific regions of interest (ROIs) of 250 × 250 μm were defined, and z-stacks were processed using the ′Sum Sliceś projection function to generate a single integrated image for intensity analysis. Quantification of EV-markers fluorescence intensity was performed from confocal immunofluorescence z-stack images using ImageJ as described above, using ROIs of 250 × 150 μm.

Cell density of IBA1 positive cells was assessed using the Cell Counter plugin, in ‘Max Intensity’ projections, using ROIs of 250 × 250 μm. To account for regions in which no cells were detected within the selected ROIs, a pseudocount of 1 was added to all raw cell counts prior to normalization. To correct for variations in section thickness, cell counts were normalized to both section depth and ROI area.

To account for inter- and intra-batch variability in reactivity marker staining, fluorescence intensity was quantified as described above, in regions close and away from the tumor, within the same section and calculations were done based on the Mean of two sections. Mean fluorescence intensity was measured as described above in matched ROIs, using ImageJ and values from areas close to the tumor were normalized to those from tumor-distal regions. Data were expressed as fold change ratios (tumor-close/tumor-away) to enable comparison across different experimental batches.

For human peritumoral tissue, two sections per sample were analyzed as described above, using a 250 × 250 µm ROI positioned within the peritumoral area in close proximity to the tumor as indicated by the operating surgeon. As a control for fold-change calculations, regions distal to the tumor were used. Control measurements were obtained from five different patients, with two sections analyzed per patient, and the mean of all control measurements was used to establish the normalization threshold.

For RNAscope image analysis, confocal immunofluorescence z-stack images were also analyzed using ImageJ. The channels were split, and GFP+ cell areas were manually defined in the GFP channel, marked as ROIs, and added to the ROI Manager. To ensure reproducibility, three single optical planes per image were selected, evenly distributed across the thickness of the z-stack. The defined ROIs were then applied to the channel corresponding to the RNAscope probe signal, and the area covered by the RNAscope probe was measured following background subtraction and threshold adjustment. All image processing parameters were maintained consistently across samples to ensure reproducibility.

Statistical analyses were performed using GraphPad Prism. Biological replicates (n) were defined as individual hc-assembloids, with group sizes represented by individual data points in each dot plot, clearly defined in figure legends. No data were excluded from the analysis, unless identified as significant outliers by a predefined outlier analysis method. Data normality was assessed using the Shapiro-Wilk test, with a significance threshold of p < 0.05 indicating non-normal distribution. Statistical comparisons were performed using the appropriate two tailed parametric or non-parametric tests based on data distribution and variance assumptions, including ordinary or Welch’s t-tests where applicable.

## SUPPLEMENTAL INFORMATION

Supplemental information can be found online at:

## AKNOWLEDGEMENTS

We are sincerely thankful to the entire Neurogenesis and Regeneration group members for their scientific inputs and discussions. We sincerely thank Dr. Judith Fischer-Sternjak for her valuable insightful contribution to the single-cell analysis. We are particularly grateful to Prof. Dr. Magdalena Götz for critical reading of the manuscript. We further acknowledge the support of the following core facilities: the Bioimaging Core Facility at BMC, the Core Facility Flow Cytometry (CFFlowCyt) at BMC and the Core Facility Pathology and Tissue Analytics at Helmholtz Zentrum München.

## Supplementary Figures

**Figure S1. Cellular heterogeneity in hc-assembloids.** (a) UMAP plot depicting the detailed annotation of the distinct 36 clusters identified in intact 130 days old hc-assembloids. Clusters are color-coded and annotated according to their respective transcriptional identity. (b, c) UMAP plots depicting localization of the telencephalic dorsal (b) and ventral (c) expression score. (d-g) Diffusion maps of astroglial clusters 9, 10,11,12, 13 (5118 cells) depicting the expression of astrocyte maturation score (d), astrocyte progenitor score (e), ependymal score (f) and astrocyte homeostatic score (g). (h) Diffusion map showing the distribution of clusters 9, 10, 11, 12, 13. (i-o) Innate microglia population in hc-assembloids. (i) Dot plot showing expression of the microglial marker genes *AIF1* and *PTPRC* in different cellular clusters. Dot size represents the fraction of cells within each cluster and color intensity indicates mean expression in group. (j) Scheme of the experimental procedure used to assess CD45 antigen expression in hc-assembloid cells with flow cytometry. (k) Dot plot depicting the fraction of CD45-positive cells in hc-assembloids. Data are presented as Mean ± interquartile range (IQR). Each dot represents a single hc-assembloid derived from the same batch. (l) Representative histogram depicting intensity of CD45 signal on hPBMCs and hc-assembloids derived microglia. (m) Dot plot showing the median fluorescence intensity of CD45 FACS signal detected on hPBMCs and hc-assembloids derived microglia. Data are presented as Mean ± standard error of the mean (SEM). Each dot represents a different donor for hPBMC and a different hc-assembloid derived from the same batch for hc-assembloids. Normality was assessed using the Shapiro-Wilk test, followed by two-tailed Welch’s t-test. (n) Dot plot showing expression of microglial sensome genes across cellular clusters. Dot size represents the fraction of cells within each cluster and color intensity indicates mean expression in group. Abbreviations: hc-human cerebral, hPBMCs-human Peripheral Mononuclear Cells, ass-assembloids, tr-transplanted

**Figure S2. Establishment of the GB–hc-assembloid model.** (a) UMAP plot depicting GBM14NSC and LN229 cells prior to transplantation (GB-C, 34256 cells). (b) UMAP plot indicating subclustering of cancer cells before transplantation showing 19 distinct clusters. (c) UMAP plot showing expression of cell cycle score across cancer cell clusters. (d) Bar plot displaying proportion of GBM14NSC and LN229 cells in different GB clusters. (e) Dot plot demonstrating GB cellular state signatures across GB-C clusters. MES1, MES2, NPC1, NPC2, AC, and OPC scores were calculated for each cluster based on Neftel et al.^45^. Dot size represents the fraction of cells within each cluster and color intensity indicates mean score in group. Color coding above the plot visualizes contribution of GBM14NSC and LN229 cells to cluster. (f-k) Micrographs depicting GFP-positive GBM14NSC (f-h) and LN229 (i-k) cells following transplantation. Scale bars: 100 μm. (l) UMAP plot visualizing the distribution of 119938 cells from all experimental groups across the dataset. (m) Dot plot showing expression score for the GFP transgene and OS9 in experimental conditions for identification of cancer cells. Dot size represents the fraction of cells within each cluster and color intensity indicates mean expression in group. (n) UMAP plot depicting 36 distinct clusters derived from hc-assembloids of all experimental conditions. Clusters are color-coded and annotated according to their respective transcriptional identity. (o) UMAP plot showing cell cycle score across the intact hc-assembloid dataset. Abbreviations: GB-C - Glioblastoma cells in culture, GB-A - Glioblastoma cells in assembloids, MES-Mesenchymal, NPC-Neural Progenitor Cells, AC-Astrocyte, OPC-Oligodendrocyte Progenitor Cells, DPT-days post transplantation, WPT-weeks post transplantation, tr-transplanted, GFP-Gren Fluorescent Protein, GB-Glioblastoma

**Figure S3. Molecular and biomechanical features of GB bulks in the hc-assembloid model.** (a) UMAP plot depicting distribution of GBM14NSC and LN229 cells before (-C) and after (-A) transplantation revealing distinct shifts in cellular states (45872 cells). (b, c) UMAP plot depicting distinct GB-C and GB-A clusters for LN229 cells (b) and GBM14NSC (c) (9180 LN229 and 36692 GBM14NSC cells). (d) Violin plot showing expression levels of *NR4A1* across LN229 clusters. (e, f) Micrographs depicting RNA scope signal for *NR4A1* RNA in LN229-C (e) and LN229-A (f) cancer cells. (g) Dot plot showing the proportion of the GFP-positive area covered by *NR4A1* RNA signal in GB-C and GB-A condition. (h) Violin plot showing expression levels of *GRID2* across GBM14NSC clusters. (I, j) Micrographs depicting RNA scope signal for *GRID2* RNA in GBM14NSC -C (i) and GBM14NSC -A (j) cancer cells. (k) Dot plot showing the proportion of the GFP-positive area covered by *GRID2* RNA signal in GB-C and GB-A condition. In panels (g, k) data are shown as mean ± SD. Each dot represents biologically independent samples (passages for GB-C and transplanted hc-assembloids from two different batches for GB-A). Normality was assessed using the Shapiro-Wilk test, followed by an unpaired two-tailed t-test. (l-n) Assessment of media dependent effects in GBM14NSC cells. (l) UMAP plot presenting GBM14NSC from three conditions, GBM14NSC-GB medium, GBM14NSC-IDM+A medium, GBM14NSC-A (65733 cells). (m) UMAP plot depicting the overlap of all experimental samples in the dataset from panel (l). (n) Heatmap showing batch effect metrics per sample compared to GBM14NSC cells in IDM+A medium. (o) Illustration of transplanted hc-assembloids showing the positioning of the magnetic ferrofluidic droplets (magenta) in tumor-free (blue) and tumor (green) areas. (p) Micrograph showing magnetic ferrofluidic droplets in GB transplanted 130-day-old hc-assembloid. (p′) is a magnified image of a tumor GFP-positive area with ferrofluidic droplets (magenta). Cell membranes were stained with CellBrite® Steady 550 Membrane dye (magenta), and tumor bulks are GFP-positive (green). (q) Dot plot depicting elastic modulus (kPa) in tumor bulks and tumor-free areas. The data are presented as the mean ± SEM for multiple areas within the same hc-assembloid. The mean for both tumor and tumor-free areas is represented by a dashed line, while the SEM is indicated by a shaded box. Scale bars: 100 μm in l, 50 μm in e, é,g, ǵ, ĺ. Abbreviations: -C-culture, -A-assembloid, GB-Glioblastoma, GFP-Green Fluorescent Protein

**Figure S4. Integration of iPSC derived neural progenitor cells (NPCs) into the hc-assembloid model.** (a) Schematic representation of the stepwise differentiation of human iPSC-derived neural progenitor cells (i-NPCs) following the protocol described by Zink et al.^68^. (b) Phase-contrast image of passage 4 (P4) i-NPCs generated from the RFP-expressing iPSC line HMGU1 (HMGU1-RFP). (c, c′′) Representative micrographs depicting expression of Pax6 (progenitor marker) and Tuj1 (neural lineage marker) in i-NPCs of P4. (d) Schematic representation of the transplantation workflow, illustrating the introduction of 20.000 RFP-positive i-NPCs into hc-assembloids. (e) Micrograph depicting RFP-positive i-NPCs, demonstrating successful engraftment of i-NPCs within hc-assembloids 3,5 weeks post-transplantation. (f-f′′′) Micrographs depicting expression of neuronal marker MAP2 and proliferation marker Ki67 in transplanted RFP-positive i-NPCs, 3,5 weeks post-transplantation. Panels (f’, f’’) present magnified images of the white boxed area in panel f. (g, ǵ, h, h′) Micrographs of i-NPC transplanted hc-assembloids showing expression of astrcytic markers GFAP (g, ǵ, h, h′), LGALS3 (g,ǵ) and CD44 (h, h′) in the transplantation site. (ǵ, h′) panels are magnified images from the white boxed areas g, h. (i-l) Micrographs demonstrating spatial pattern of glial reactivity upon transplantation of i-NPCs in areas close (i,k) and away (j, l) from the transplantation site. (m, n) Dot plots presenting the mean fluorescence signal intensity of GFAP-(m) and LGALS-(n) positive signal in areas close and distant to NPC-transplantation site. (o) Dot plot showing the cell density of IBA-positive cells in areas close and distant to NPC-transplantation site. Paired measurements are connected with dashed lines. Single dots represent single transplanted hc-assembloids obtained from 4 independent batches. Data normality was assessed with Shapiro-Wilk normality test in all groups. Statistical comparisons were performed using a two-tailed paired t-test for panels (m, n) and a two-tailed Wilcoxon matched-pairs signed-rank test for panel (o). Scale bar: 100 μm in e, f-f’’, g, h; 50 μm in b, ǵ, h′, i-l and 20μm in c, c′. Abbreviations: iPSCs-Induced Pluripotent Stem Cells, EBs-embryoid bodies, RFP-Red Fluorescent Protein, RFP-Red Fluorescent Protein, NPC-Neural progenitor cells

**Figure S5. Microglial reactivity in the peritumoral area.** (a-é) Micrographs depicting distribution (a, b, c, d, e) and morphology (á, b′, c′, d′, é) of IBA1-positive microglia in the intact (a, á), media transplanted (b, b′), i-NPC-(c, c′), LN229-(d, d′) and GBM14NSC-transplanted (e, é) hc-assembloids. Images in á, b′, c′, d′, é are magnifications of white boxed areas in a, b, c, d, e respectively. (f, g, I, j) Micrographs showing accumulation of IBA-positive cells in areas proximal (f, i) and distal (g, j) to GB revealing the spatial pattern of microglial reactivity in GB14NSC-(f, g) and LN229-(i, j) transplanted hc-assembloids. (h, k) Dot plots showing cell density of IBA-positive cells in areas close and distant from GBM14NSC-(h) and LN229 (k) GB bulk. Single dots represent single transplanted hc-assembloids obtained from ≥7 independent batches. Data normality was assessed with Shapiro-Wilk normality test in all groups. Statistical comparisons were performed using a two-tailed paired t-test. (I, m) Micrographs indicating the absence of microglia cells in intact (l) and GBM14NSC-transplanted (m) hc-organoids. (n, o) Micrographs showing GFAP and LGALS3 immunoreactivity in areas proximal (n) and distal (o) to GB in GBM14NSC-transplanted hc-organoids. (p, q) Dot plots depicting GFAP (p) and LGALS3 (q) mean fluorescence signal intensity in areas close and distant from the GB bulk. Paired measurements are indicated by dashed lines. Single dots represent single transplanted hc-organoids obtained from one batch. Data normality was assessed in both groups with Shapiro-Wilk normality test and comparisons made using two-tailed paired t-test. (r) UMAP plot depicting color coded separation of cancer cells and hc-assembloid derived cells (37428 cells) (s) UMAP plot presenting cell annotation by regional identity of cells (dorsal, ventral and cells with no unique identity (N/A)). (t) Heatmap showing the strength of inferred cell-cell communication between ventral assembloid cells, dorsal assembloid cells, and cancer cells. Scale bars: 100 μm in a, b, c, d, e and 50 μm in á, b′, c′, d′, é, f, g, I, j and l-o. Abbreviations: GB-Glioblastoma, hc-human cerebral, ass-assembloid, NPCs-Neural Progenitor Cells, tr-transplanted, Or-organoids

**Figure S6. Gene Expression Signatures of Glial Cells Induced by GB Transplantation.**(a) UMAP plot depicting distribution of hc-assembloid derived cells across experimental conditions (73585 cells). (b) UMAP plot showing cellular heterogeneity of hc-assembloid derived cells of media-, GBM14NSC- and LN229-transplanted hc-assembloids. (c) Boxplot illustrating batch effect metrics per cell label across hc-assembloid conditions using intact hc-assembloid condition as background control. (d, e) Bar plots showing the top 20 significantly enriched Gene Ontology (GO) Biological Process terms (p < 0.05), ranked by fold enrichment, for astroglial cluster 1 (d) and cluster 3 (e), enriched in transplanted hc-assembloids. Color intensity indicates the number of genes associated with each GO term (term size). (f, g) Microglia reactivity on day 132 of hc-assembloids. Dot plots showing expression of microglial homeostatic (f) and reactive (g) genes in intact, media-transplanted and GB-transplanted hc-assembloids. Dot size represents the fraction of cells within each cluster and color intensity indicates mean expression in group. (h, j, l) Spatial maps depicting the spatially smoothed general malignancy score in the infiltration zone of the human IDH-wildtype glioblastoma samples derived from the publicly available spatial transcriptomic dataset of Greenwald et al. ^80^ for each section. (i, k, m) Spatial maps depicting the spatially smoothed score of the 15-gene common reactive astrocyte signature projected onto the corresponding sections. Each spot is color-coded according to the adjacent scale. Abbreviations: tr-transplanted, hc-human cerebral, GB-Glioblastoma, inf-infiltrative zone

**Figure S7. Differentiation, phenotypic Characterization, and integration of i-monocytes into the hc-assembloid model.** (a) Schematic overview of i-monocyte differentiation from hiPSCs, following a modified version of the Cao et al. protocol ^90^. Phase-contrast image displays the characteristic morphology of i-monocytes obtained with this differentiation protocol. (b) Dot plot depicting the proportion of CD14+ and CD16+ cells following i-monocyte differentiation from three different hiPSC lines (HMGU1, HMGU1-RFP, and HMGU12). Data are shown as mean ± SEM. Each dot represents an independent differentiation starting from different hiPSC lines as indicated. (c, e, g) Scheme illustrating the experimental protocol for immunostaining-based characterization of i-monocytes (c), assessment of their phagocytic capacity using pHrodo™ BioParticles™ Conjugates for Phagocytosis (e), and i-monocyte transplantation into hc-assembloids (g). (d-f′′) Representative micrographs depicting the expression of mononuclear cell lineage markers IBA1 and CD14 (d-d′′) and capacity of i-mono to uptake the pHrodo beads (f-f′′). Images d′, d′′, f′ and f’′′ present a close-up of the white-boxed region in panels d and f, respectively. (h-í′′) Representative micrographs showing integration of CD14+ and MPEG1+ i-monocytes in intact (h-h′′′) and hc-assembloids transplanted with GFP-labelled GB cells (i-í′′) 7 days after i-monocyte inoculation. Images in h’-h’’’, and i’-i’’’ present a close-up of the white-boxed region in panels h and i, respectively. Scale bars: 100 μm in h-i; 50 μm in a, d, f, h′,h′′,h′′′,í, í′,í′′; 20 μm in d′, d′′, f′, f’′′. Abbreviations: hiPSCs-Human Induced Pluripotent Stem Cells, 2wpt-2 weeks post transplantation, 3wpt-3 weeks post transplantation, PHRODO-pHrodo™ BioParticles™ Conjugates for Phagocytosis, tr-transplanted.

**Video S1.** Video presenting GFP-positive tumor bulb (green) within a scilce of a LN229 transplanted hc-assembloid where GFP-positive cells interact with resident cells from the peritumoral area (magenta).

**Video S2.** Video of a sliced LN229-transplanted hc-assembloid showing the active invasion of GFP-positive GB cells (green) into the surrounding tumor-free tissue (magenta).

**Video S3.** Long-term live imaging of GB invasion into brain tissue in an air-liquid interphase cultured tumor hc-assembloid slice with GFP-positive GBM14NSC cells (green) and nuclei labeled with NucSpot 650, predominantly representing cells of the non-tumor brain tissue (red). At each time point, a 71-µm z-stack was acquired with a 1-µm z-step and rendered as a maximum-intensity projection.

**Video S4.** Video depicting the deformation of a ferrofluidic droplet (red) within a tumor mass (green), upon application of an applied magnetic stress in a transplanted hc-assembloid.

**Video S5.** XYZ stack of a cleared intact hc-assembloid a week after inoculation with i-monocytes (magenta), stained with GFAP (white).

**Video S6.** XYZ stack of a cleared transplanted hc-assembloid a week after inoculation with i-monocytes (magenta) and GFP (green) stained with GFAP (white).

