## Supplementary material for "Human cerebral assembloids as a model to study the interaction of glioblastoma with resident glial cells": Suppl Figures

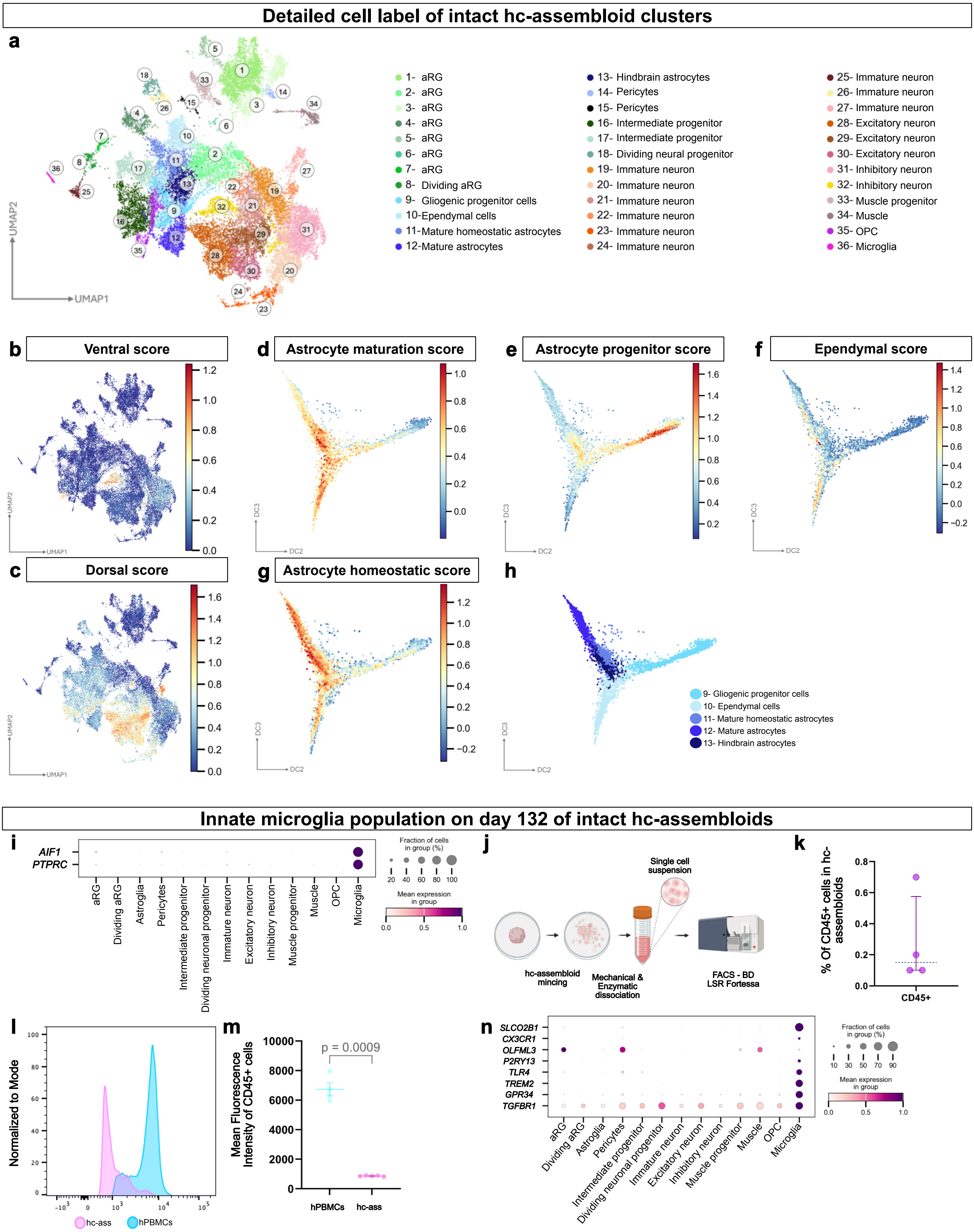


Kalpazidou S., Moysidou-Tsiorva P., et al, Figure S1

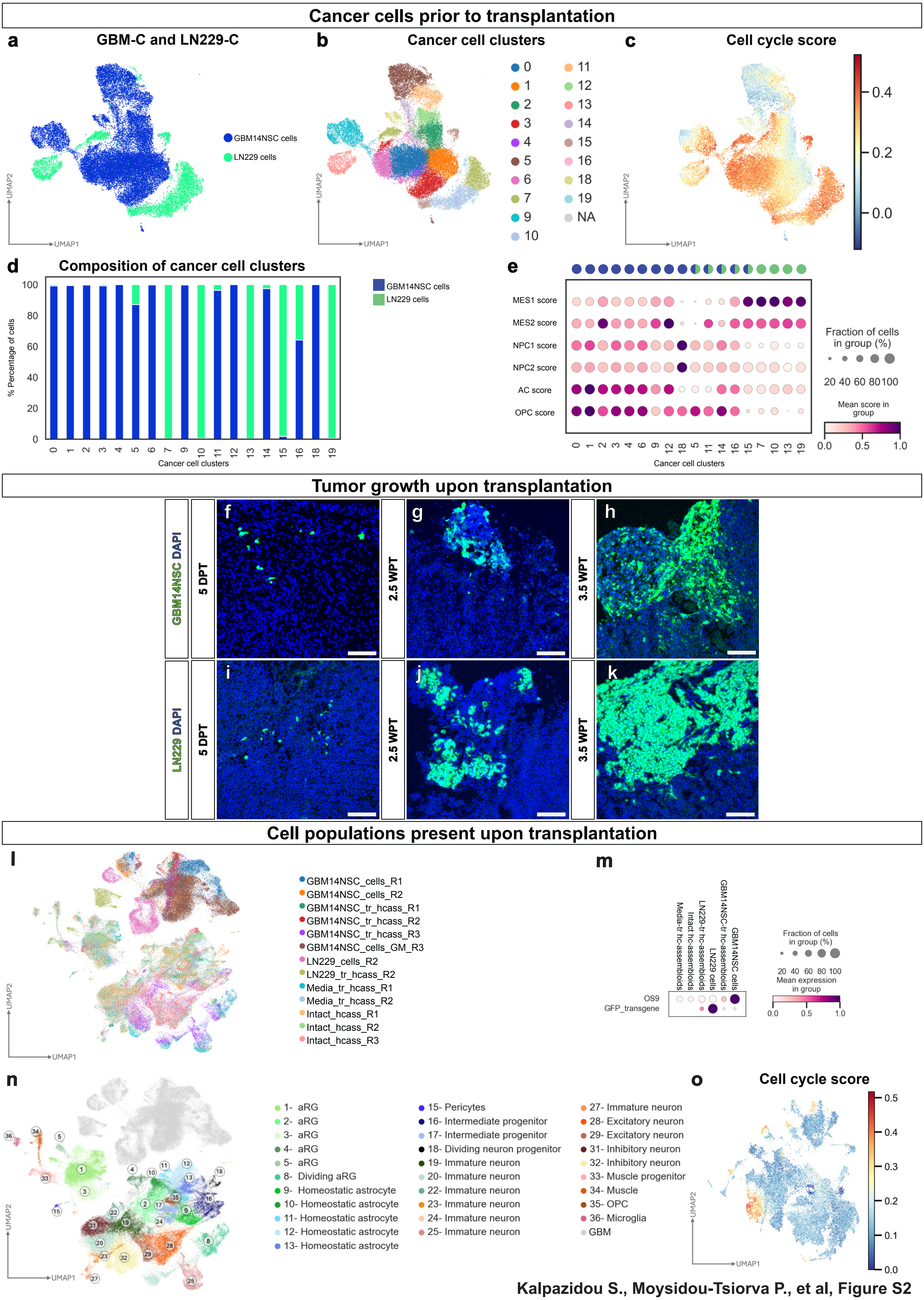


Kalpazidou S., Moysidou-Tsiorva P., et al, Figure S2

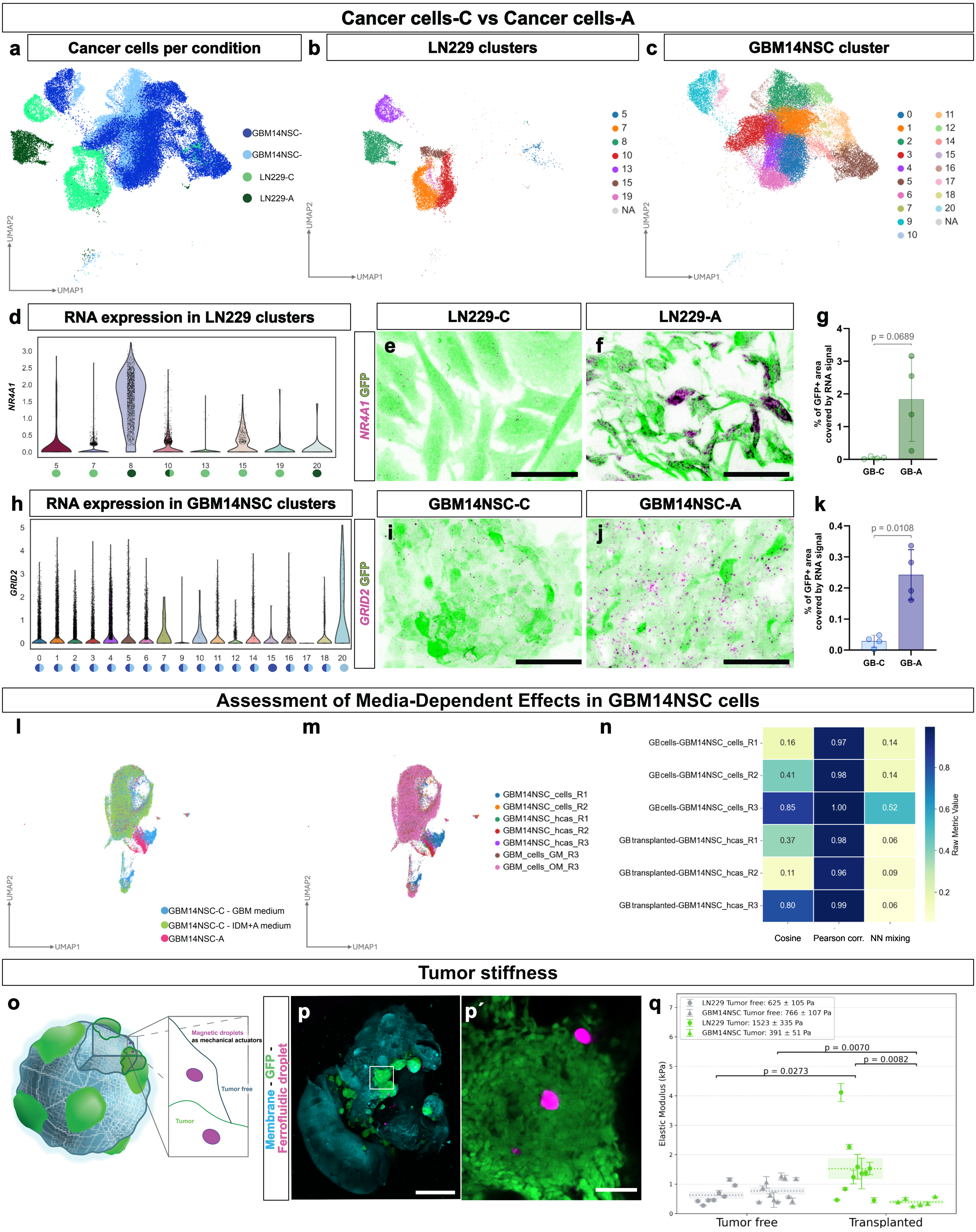


Kalpazidou S., Moysidou-Tsiorva P., et al, Figure S3

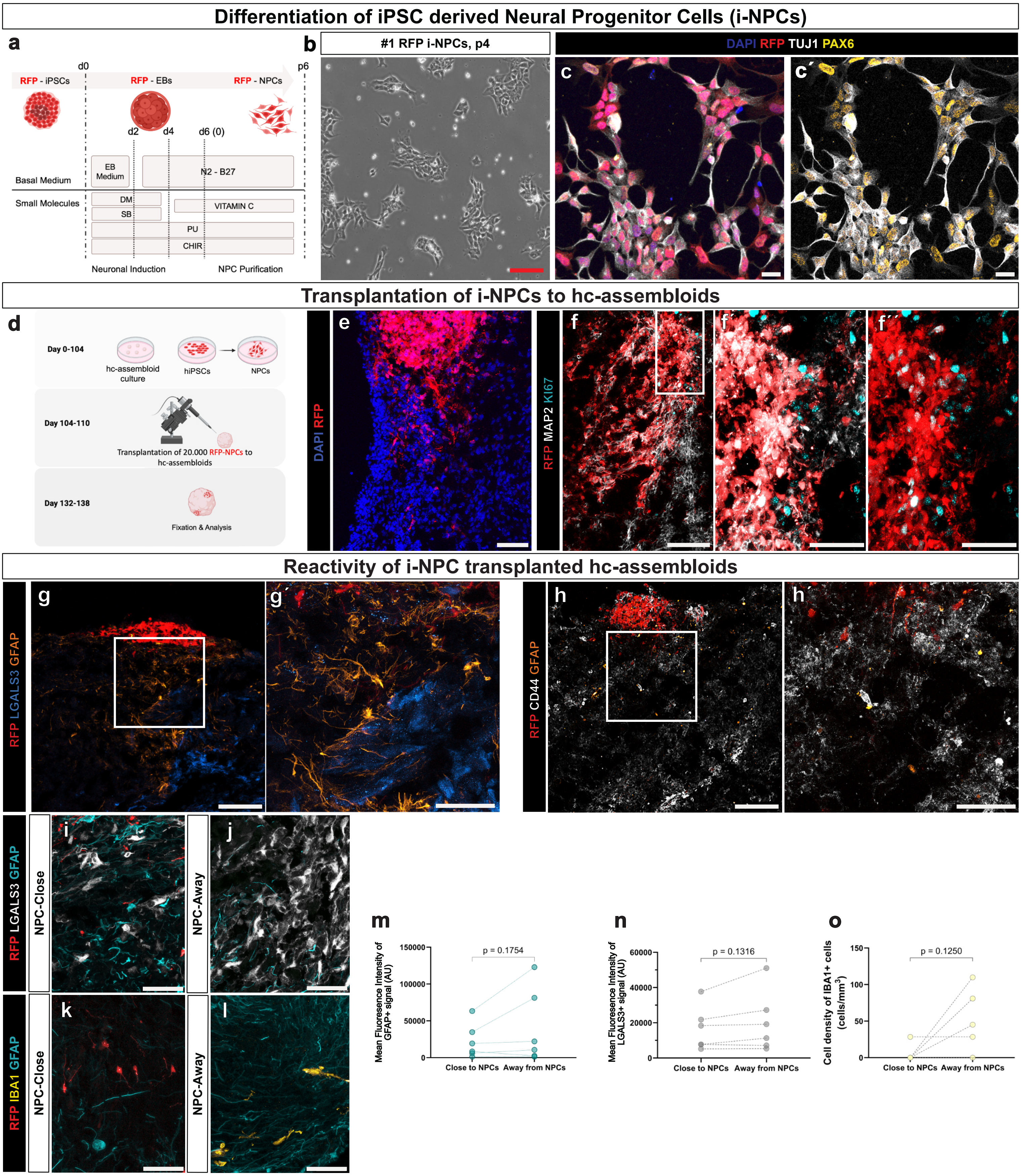


Kalpazidou S., Moysidou-Tsiorva P. et al., Figure S4

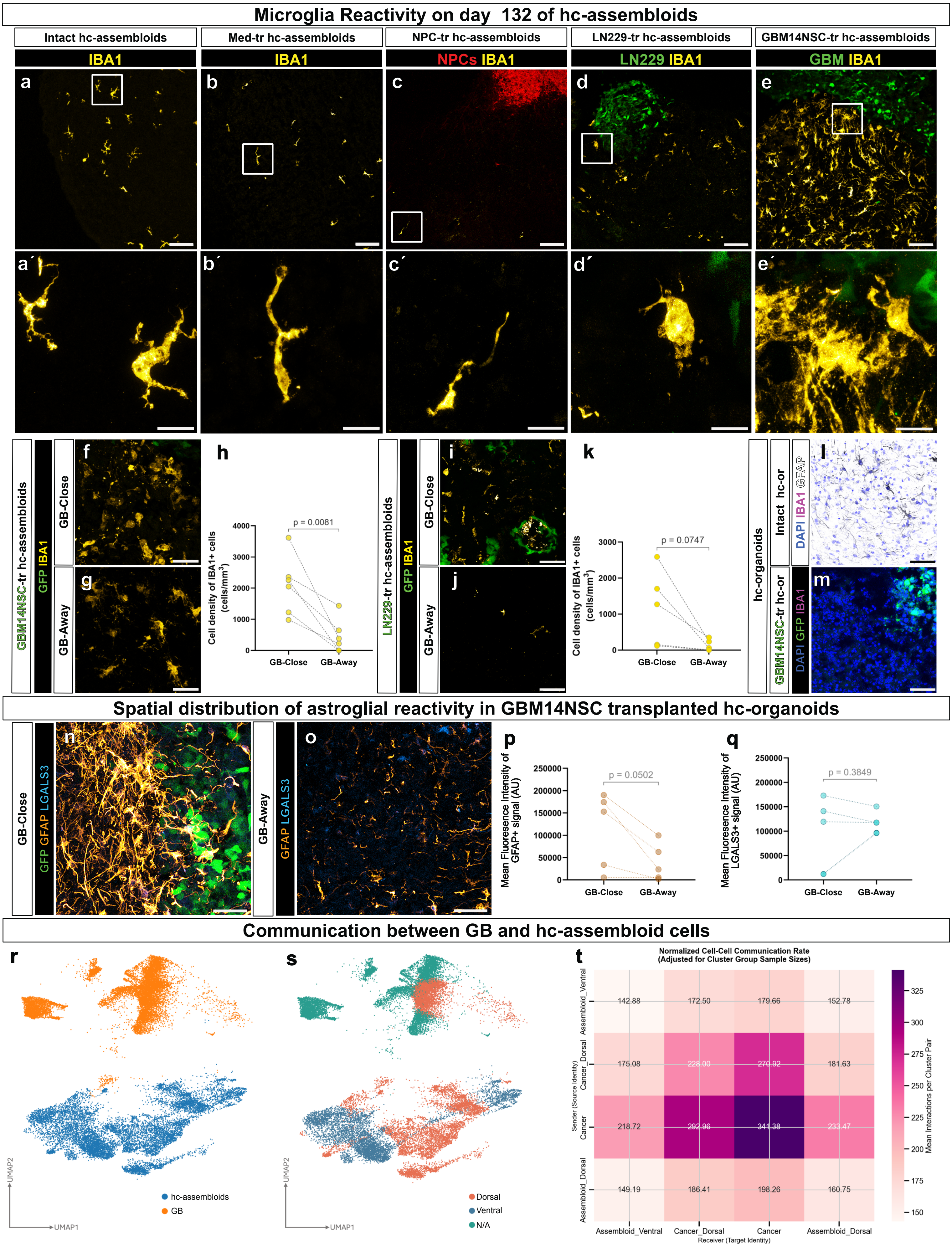


Kalpazidou S., Moysidou-Tsiorva P., et al, Figure S5

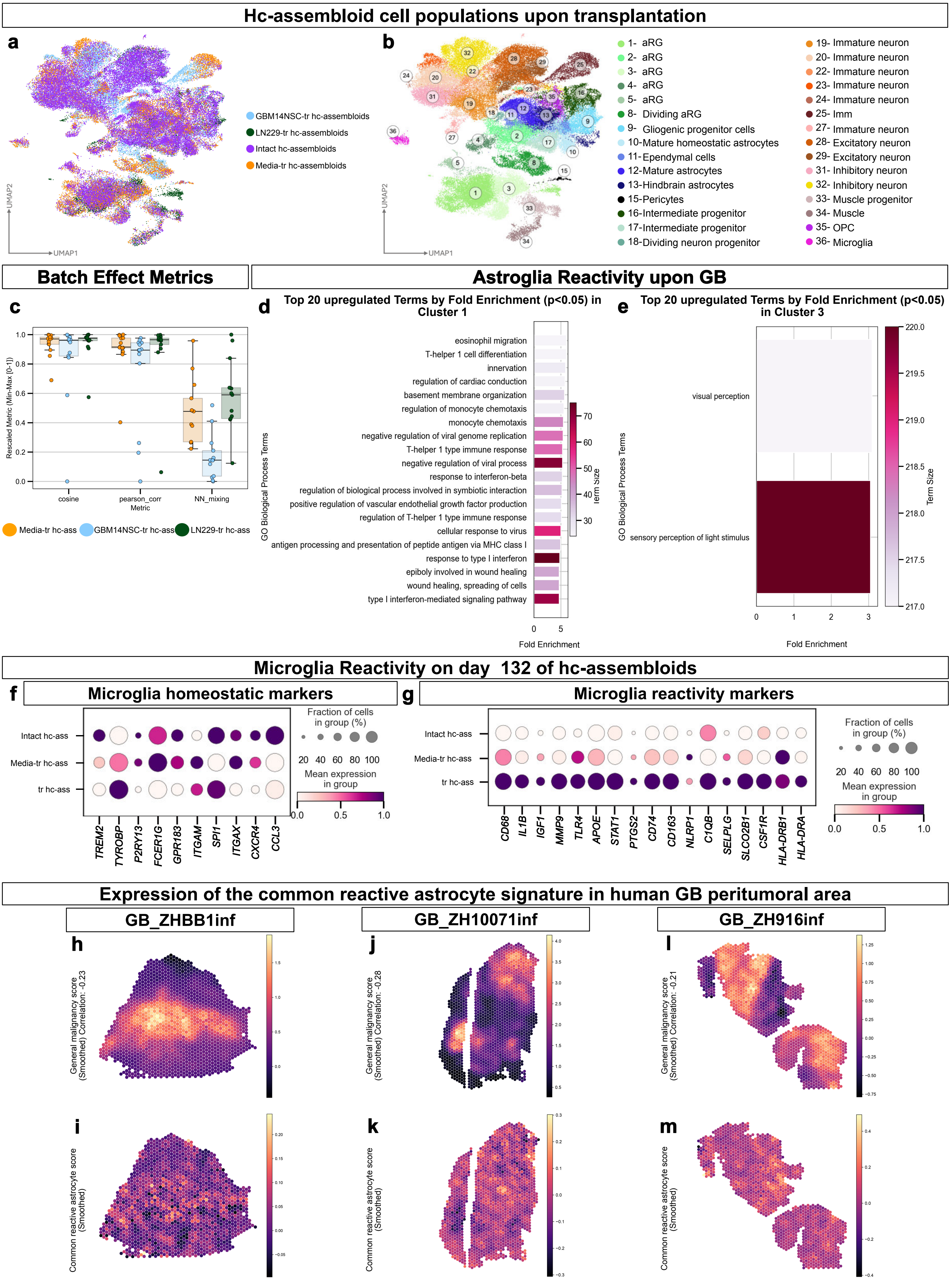


Kalpazidou S., Moysidou-Tsiorva P. et al, Figure S6

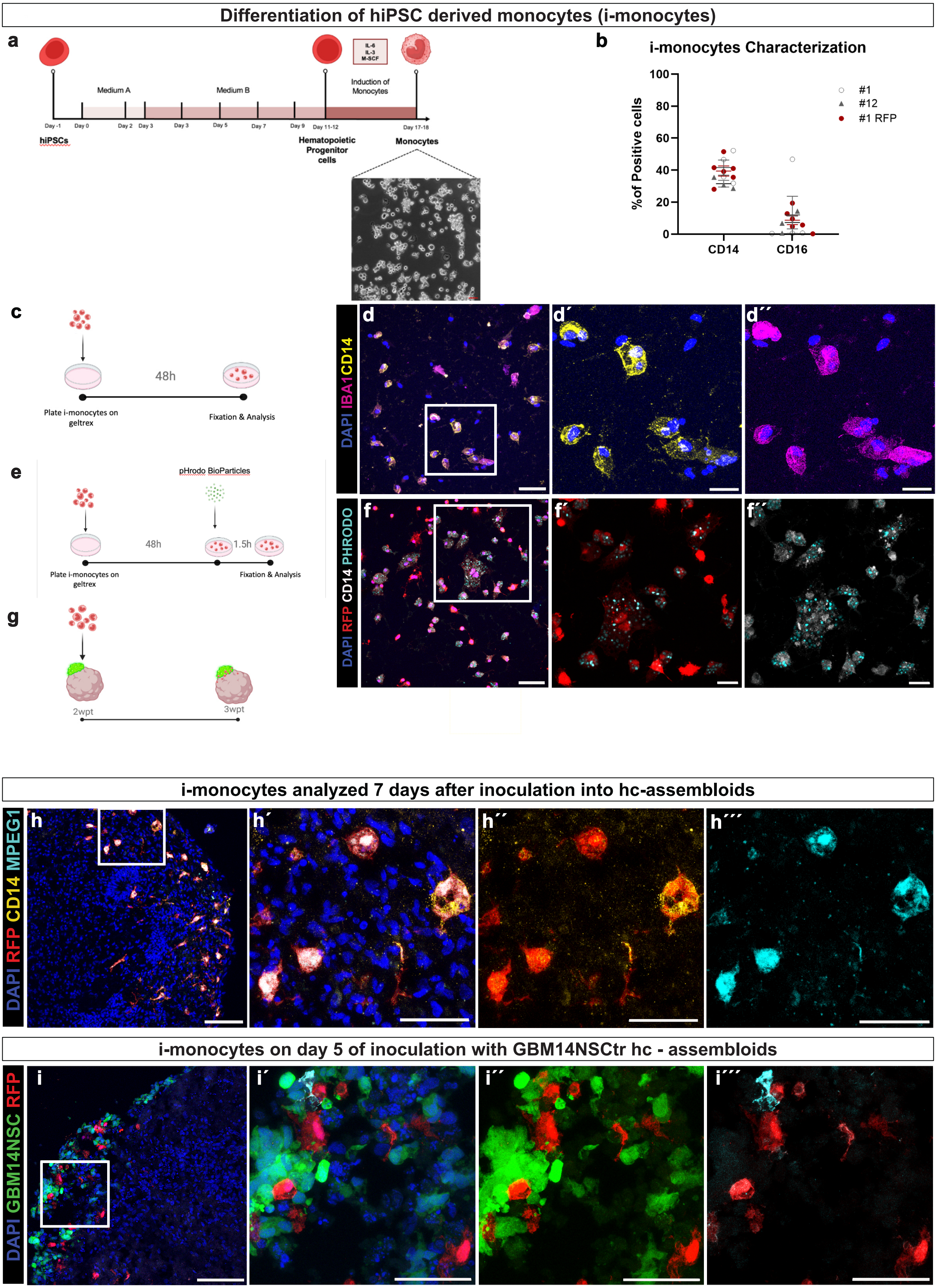


Kalpazidou S., Moysidou-Tsiorva P. et al., Figure S7
